# Exploring the potential of using simulation games for engaging with sheep farmers about lameness recognition

**DOI:** 10.1101/2022.10.26.513828

**Authors:** Matt L. Jones, Maxwell S. Barnish, Robert R. Hughes, Aimee Murray, Omid Mansour, Tiziana Loni, Holly Vickery, Myfanwy Lloyd Evans, Laura Green, Nervo Verdezoto

## Abstract

**Introduction:** Computer simulation games are increasingly being used in agriculture as a promising tool to study, support and influence real-life farming practices. We explored the potential of using simulation games to engage with sheep farmers on the ongoing challenge of reducing lameness. Working with UK stakeholders, we developed a game in which players are challenged with identifying all the lame sheep in a simulated flock. Here, we evaluate the game’s potential to act as a tool for to help assess, train and understand farmers’ ability to recognise the early signs of lameness.

**Methods:** Participants in the UK were invited to play the game in an online study, sharing with us their in-game scores alongside information relating to their real-life farming experience, how they played the game, and feedback on the game. Mixed methods were used to analyse this information in order to evaluate the game. Quantitative analyses consisted of linear modelling to test for statistical relationships between participants’ in-game recall (% of the total number of lame sheep that were marked as lame), and the additional information they provided. Qualitative analyses of participants’ feedback on the game consisted of thematic analysis and a Likert Scale questionnaire to contextualise the quantitative results and identify additional insights from the study.

**Results:** Quantitative analyses identified no relationships between participants’ (n = 63) recall scores and their real life farming experience, or the lameness signs they looked for when playing the game. The only relationship identified was a relationship between participants’ recall score and time spent playing the game. Qualitative analyses identified that participants did not find the game sufficiently realistic or engaging, though several enjoyed playing it and saw potential for future development. Qualitative analyses also identified several interesting and less-expected insights about real-life lameness recognition practices that participants shared after playing the game.

**Discussion:** Simulation games have potential as a tool in livestock husbandry education and research, but achieving the desired levels of realism and/or engagingness may be an obstacle to realising this. Future research should explore this potential further, aided by larger budgets and closer collaboration with farmers, stockpeople and veterinarians.

## Background

Lameness is a change in animal gait that has various underlying causes, but is typically caused by bacterial infections of the hoof and foot (especially scald and foot rot) in farmed sheep, goats and cattle (Kaler et al. 2019). As a macro-level manifestation of microbial ailments, the first diagnosis of lameness can typically made by farmers after visual observation of their livestock walking. Despite this, lameness is still a major burden on livestock farming, with some evidence that this is partly because farmers differ in their ability to recognise lameness, especially in its early stages (Whay et al. 2003; Green and Clifton 2018). In UK sheep farming, lameness is estimated to cost farmers between £3.90 and £6.30 per ewe per year (Winter and Green 2017), and the industry as a whole £28-80 million per year (Nieuwhof and Bishop 2005; Wassink et al. 2010). As well as economic costs associated with veterinary expenses and livestock productivity losses, lameness also constitutes a substantial animal welfare (FAWC 2011; Nalon and Stevenson 2019) and antibiotic stewardship problem (Davies et al. 2017), making it a priority issue for the sheep farming industry to address. In 2011, the Farm Animal Welfare Council (FAWC) challenged UK sheep farmers to reduce the average prevalence of lameness on UK sheep farms to less than 5% by 2016 and less 2% by 2021 - targets that were, at the time, considered achievable using evidence-based techniques (FAWC 2011). Whilst the initial 5% target appears to have been met - with a well-randomised study estimating the mean flock prevalence of lameness in the UK to be 3.5% (ewes) in 2013 (Winter et al. 2015) - there are signs that progress may have since stalled. The most recent (though non-randomised) study estimated a mean flock prevalence of lameness (ewes) of 3.2% in the 2018-2019 period, suggesting that farmers were not on track to reach the 2021 2% target (Best et al. 2021). Furthermore, there are indications of limited uptake and farmer scepticism towards some of the lameness-reduction techniques recommended by the FAWC (Best et al. 2020, 2021), and that the numbers of farmers practicing key effective treatments may be reducing over time (Prosser, Purdy, and Green 2019). Collectively, these observations suggest that new approaches might be needed to facilitate knowledge exchange between farmers and other interested parties to reduce lameness in the UK.

One new strategy to facilitate knowledge exchange between farmers and non-farmers that has recently been explored in agricultural education and research is the use of game-based approaches to facilitate innovation, participation and multiple stakeholders perspectives (Hernandez-Aguilera et al. 2020; Berthet et al. 2016). The progress of information and communication technology (ICT) has led to the development of farm-based computer and video games worldwide that have actively engaged players in virtual farming environments (Sutherland 2020). Indeed, computer-mediated virtual agricultural environments are well-established as mass-appeal simulation video games such as FarmVille and Farming Simulator, which serve as forms of entertainment for non-farmers and farmers alike (Lane 2018). However, more recently, virtual environments have begun to be used as pedagogic and research tools for engaging with farmers in order to address serious, real-world issues. Most commonly, researchers have explored the use of virtual environments for educational purposes, having benefits such as making agricultural training more logistically feasible, affordable and accessible (Barber 2016). Several projects have developed and explored the potential of games of this sort - including developing games for teaching crop cultivation and livestock breeding skills (Yoo and Kim 2014; Szilágyi et al. 2017), developing more all-encompassing agricultural training games (GATES 2019; Fountas, Spyros et al. 2019), and exploring the potential of virtual reality-assisted agricultural training (Barber 2016). Virtual agricultural environments may also serve less obvious knowledge exchange purposes; for example, to encourage the adoption of precision agriculture technologies (Pavlenko et al. 2021); to exchange knowledge and perspectives on farm design among farmers, researchers and advisors (Moojen et al. 2022); to facilitate information sharing among farmers and with non-farmer stakeholders dealing with agricultural issues (Hernandez-Aguilera et al. 2020; Nuritha, Widartha, and Bukhori 2017). The idea of using virtual environments as tools for engaging with farmers is thus being taken increasingly seriously; representing a new, innovative, participatory, and even fun approach to understanding and addressing the real-world challenges of modern agriculture.

Here, we explore the potential of using computer-based gaming as an innovative approach to engage with UK sheep farmers and other stakeholders on the issue of the early recognition of the signs of lameness. Sheep lameness can be graded according to increasing severity of change in gait, and sheep farmers recognise different severities of lameness innately (Kaler and George 2011). Farmers that report that they recognise, catch and treat the first mildly lame sheep in a group experience lower prevalences of lameness compared to farmers who wait until sheep are more severely lame before they catch them (Kaler and Green 2008; Winter et al. 2015). Following a human-centered design approach, we developed a game (The Lameness Game) that is intended to support lameness reduction by serving as a tool to help assess, train and understand farmers’ ability to recognise the early signs of lameness. We evaluated our game through an online evaluation study with participants playing and giving expert feedback on our prototype game, reporting our analysis of their in-game performance and feedback in order to assess the games’ potential.

## Materials and methods

### Description of The Lameness Game

Our game was a single-player, casual simulation game in which players were set the goal of identifying all of the lame sheep in a virtual flock in the shortest time possible (Figure 1). During gameplay, the displayed environment resembles a farm field which is occupied by virtual sheep programmed to spend most their time grazing (~73% of the time) or standing (~23.5% of the time), but that occasionally walked (~3.5% of the time). These parameters were intended to be somewhat reflective of estimated real-life ovine activity budgets whereby walking constitutes a minority (~2-4%) of the total activity (Kaler et al. 2019; Bueno and Ruckebusch 1979), whilst also providing a small (but not impractically small) window of opportunity to identify lame sheep within the time-frame of a relatively short game. Players could navigate the environment with game controls that resemble those of a simplified real-time strategy game; up-down-left-right to move the camera to move the camera across the field (WASD keyboard keys), camera rotate to change the direction of camera (Q & R keyboard keys) and zoom controls to change the field of view of the camera (trackpad/mouse scroll). At the start of the game, a ‘healthy’ or ‘lame’ status is randomly assigned to each of the 24 sheep in the flock (i.e. on average 50% of the sheep were assigned to be lame via a coin-flip style mechanism, though this was not disclosed to the player), which determines the animation used when they walk (Figure 1A). In our game, lame sheep exhibited a shortened stride on one (infected) leg, a quickened stride on the opposite leg, and a slight nodding of the head - approximating the signs of early lameness represented by Score 2 on the scale. When players identified a sheep they thought was lame, they could select it by clicking it with the left mouse button, upon which an icon appeared above the sheep’s body that the users could click to mark the sheep as lame (Figure 1B). The sheep was then marked with a purple spray and its status changed to ‘Marked as Lame’ for the purposes of the in-game scoring system. At the end of the game, users received a score for their accuracy (% of sheep marked that were actually lame) and recall (% of the total number of lame sheep that were marked as lame), some educational feedback on their performance, as well as the time remaining on the in-game clock (Figure 1C). Players were given a maximum of ten minutes to identify the lame sheep, but could choose to terminate the game and get their results early by clicking ‘Done’.

**Figure 1:**
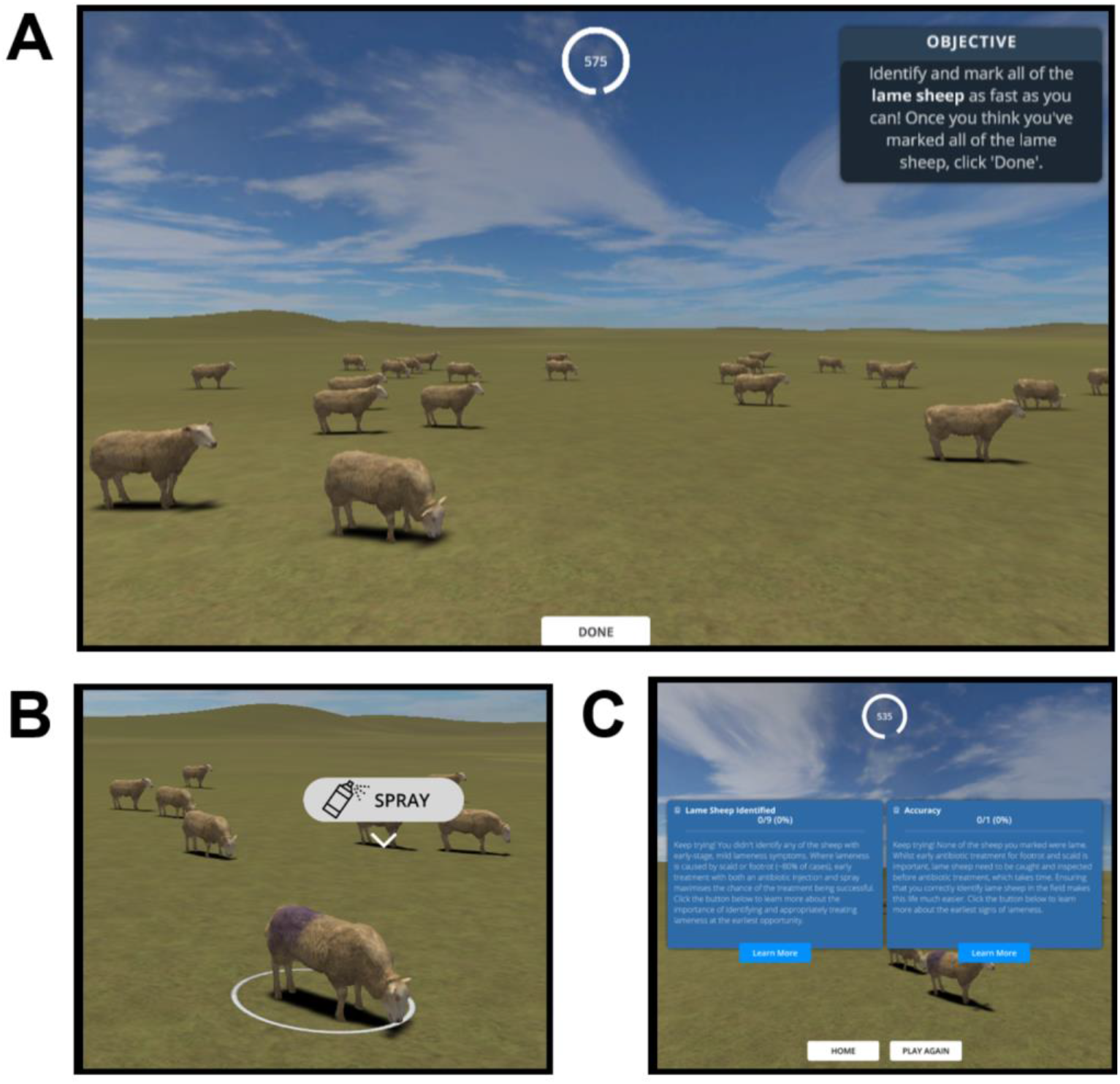
Screenshots of summarising the main features of the game. A) In the game, players are presented with a field of virtual sheep and the goal of observing them to identify those with a lame gait. B) Users can zoom in and select sheep, spraying them purple to mark them as lame C) At the end of the game (10 minute timer ends or users click ‘Done’), users are presented with scores based on how many of the sheep they marked as lame were actually lame, as well as some related educational information.

The game was developed using a human-centered design (HCD) process in which potential users (farmers, farm veterinarians and academics in the field) were involved throughout all stages of the design process (Hanington 2017), and substantially shaped the final game we evaluate here (Supplementary Material 1; Supplementary Figure 1). The final game was built using game-programming software Unity and 3D modelling software Blender (Blender Foundation 2021) in collaboration by a game programmer (OM) and 3D artist/animator (TL) using a mix of pre-made, modified and newly-created 3D models, animations and other digital assets (Red Deer 2020; Bicameral Studios 2018; Lehtonen 2017; Michsky 2021). The game runs standalone in a browser on desktop and laptops, preferably using the Google Chrome browser. A playable version of the game is available free of charge online (https://wheres-woolly.itch.io/lameness-game) and/or from the corresponding author.

### Evaluation of The Lameness Game

The game was evaluated via an online study in which those with and without agricultural experience were invited to play the game online and fill in an after-game questionnaire via the Microsoft Forms platform (Supplementary Material 2). Through the after-game questionnaire, participants shared with us their in-game scores (those presented via the screen shown in Figure 1C) alongside feedback on the game. Participants were enrolled in the study by advertising it on social media and private mailing lists (targeting groups of interest where possible e.g. sheep societies), as well as during a workshop with University of Bristol Farm Animal Discussion Group (comprising veterinary practitioners, teaching staff and researchers). Participation was incentivised by offering participants entry into a lottery to win one of three £50 vouchers for an online farm supplies shop in return for the approximately 30 minutes of participation time. This study was approved by the College of Medicine and Health research ethics committee at the University of Exeter (application number 21/01/275). To comply with ethical requirements, participants were required to read an information sheet and digitally sign a consent form before participating in the study.

#### Participant recall scores in the game

Quantitative evaluation of the game consisted of analysing the relationship between participants’ recall scores in the game and data relating to their real-life experience and how they played the game (all self-reported in the after-game questionnaire; Supplementary Material 2). Our logic was that the game could serve as a tool for training, testing or studying real-life lameness recognition practices if participants were able to translate real-life experience and skills into higher in-game recall scores. Recall was calculated and reported alongside accuracy at the end of the game (Figure 1C) and as for all other data, participants shared these scores with the research team via the after-game questionnaire.

In order to test whether participants had played the game as intended before engaging in further analysis, we first used D’agostino’s test to test for normality and skewness in participants’ recall and accuracy scores. A range of recall scores is expected to be underpinned by generally high (negatively skewed) accuracy scores (i.e. the majority of scores above >50%) if participants had successfully engaged with the goal of the game (to mark all the sheep they think are lame) without ‘cheating’ (i.e. by taking a ‘shotgun’ approach and marking all sheep as lame). High accuracy scores also gave us a first indication that our animations of lameness were at least realistic enough for participants’ to recognise as lameness.

Subject to confirming this, we then proceeded with a more quantitative analysis of participants’ recall scores; seeking to identify a feasible linear model describing what (if anything) affected participants’ recall scores (subject to them meeting the assumption of normality). In order to do this, a post-hoc power analysis was first performed to understand how complex a model we could build with the sample size (power) available. Accounting for our sample size (n = 63), assuming stringent 95% power and 5 significance thresholds, and the use of a linear model with 1 on 61 degrees of freedom (i.e. a single continuous or two-factor explanatory e.g. true-false type variable), we estimated that our study had the power to detect an approximately ‘medium-to-large sized’ effect (f2 = 0.21), *sensu* Cohen (1977). Accordingly, we tested different candidate linear models - each with a single explanatory variable describing what drove participants’ ability to identify lame sheep in the game - until a feasible model was identified. Beginning with our first hypothesis that there was a relationship between participants’ in-game scores and their real-life farming experience (‘Farming Experience’ hypothesis), we progressed through to models testing for an effect of lameness signs participants looked for during the game (‘Lameness signs looked for’ hypothesis), and finally for an effect of more idiosyncratic factors to do with user engagement (‘User engagement’ hypothesis). To choose the explanatory variable computed in each model considered, we used an exploratory data analysis approach (Tukey 1977); plotting all variables relating to the hypothesis under consideration, and then choosing the variable(s) that visually appeared to have the strongest effect on recall scores for modelling (helping to mitigate against issues caused by multiple hypothesis testing). For the ‘Farming Experience’ hypothesis, candidate variables plotted and chosen from were: whether or not the participant had experience in farming/related field (TRUE/FALSE categorical variable of 2 levels derived from Q15 in the questionnaire); the perceived annual prevalence of lameness they had experienced if they had farming experience (categorical variable of 2 levels derived from Q19 in the questionnaire); the number of years they had spent working with sheep if they had farming experience (continuous variable derived from Q19 in the questionnaire). For the ‘Lameness signs looked for’ hypothesis, the candidate variables were the 9 signs of lameness that participants told us they did or did not look for e.g. uneven posture, shortened stride on one leg when walking (TRUE/FALSE categorical variables of 2 levels derived from Q13 in the questionnaire). For the ‘User engagement’ hypothesis the candidate variables were: how many times the participant had played the game before submitting their scores (categorical variable of 5 levels derived from Q5 in the questionnaire); whether or not the participant had problems with the game’s controls (TRUE/FALSE categorical variables of 2 levels derived from Q7 in the questionnaire); observing type/how the participant observed the sheep when playing the game (categorical variable of 3 levels derived from Q10 in the questionnaire); moving type/how the participant moved around the flock when playing the game (categorical variable of 4 levels derived from Q11 in the questionnaire); whether or not the participant completed the pre-game tutorial (categorical variables of 3 levels derived from Q6 in the questionnaire); the computer set-up/pointing device the participant used (categorical variables of 3 levels derived from Q7 in the questionnaire); and the time spent playing the playing (continuous variable derived from Q2 in the questionnaire). In total, we tested four models - one for the ‘Farming experience’ hypothesis, two for the ‘Symptoms looked for’ hypothesis, and one for the ‘User engagement’ hypothesis. P-values from each of the models were Bonferroni-corrected according to the number of previous models tested, and we stopped building models once a feasible model was identified (i.e. one with a p-value < 0.05). Our null hypothesis (H0) in all models was that our measured variable(s) did not affect participants’ recall, whilst our alternative hypotheses was that the variable under consideration affected participants’ recall.

This analysis was performed in the R programming language (R Core Team 2017) implemented via RStudio (RStudio Team 2020). Exploratory plotting to identify candidate variables for linear modelling was conducted using base R functions and the *beeswarm* function of the ‘beeswarm’ package (Eklund and Trimble 2021). Given that accuracy and recall scores were percentage data, they were both arcsine square root transformed using base R functions before being subjected to statistical testing (D’agostino’s test and linear modelling). D’agostino’s test was implemented via the *agostino.test* function of the ‘moments’ package (Komsta and Novomestky 2022). Power analysis was implemented via the *pwr.f2.test* function of the ‘pwr’ package (Champely et al. 2020). Linear modelling and Bonferroni correction of p-values was performed using base R functions.

#### Feedback on the game from those with real-life farming experience

To help explain the results from the quantitative analysis of participants’ recall scores and evaluate the game more broadly, we also collected feedback on the game from participants who had real life farming experience and conducted complementary qualitative analyses. We limited this data collection and evaluation to participants who had worked in farming or a related field (i.e. those who had answered ‘Yes’ to the question ‘Have you ever worked in farming or a related field e.g. farm vet?’) because this was the intended audience of the game. These participants with real-life farming experience directly evaluated the game in two ways; by providing open-form feedback in the after-game questionnaire, and by scoring evaluation statements on a Likert scale.

Open-form feedback provided an opportunity for participants to elaborate on their thoughts about the game and suggest new potential uses of it. This feedback was analysed using inductive thematic analysis, a qualitative analytical technique that involves finding patterns in a non-numerical dataset to understand participants’ opinions, perspectives and experiences (Braun and Clarke 2006, 2021b). Thematic analysis values all participants’ perspectives without privileging the ore commonly/frequently expressed perspectives that might prioritise the quantification of patterns e.g. coding reliability approaches, underpinned by positivist approaches and quantitative methods (Braun and Clarke 2021a, 2021a). We conducted thematic analysis on free-text feedback from those who provided it (n = 19, from the total of 31 participants with real-life farming experience). Statements were coded and then reported in terms of themes, each consisting of one or multiple conceptually linked sub-themes. Supporting quotes were noted to illustrate each sub-theme. Analysis was initially conducted independently by two researchers (MSB and NVD) reading and coding all free-text feedback and identifying the initial themes. Any discrepancies (e.g. disagreements in assignment of comments to themes, comments fitting more than one theme) were initially discussed between these two researchers then an agreed analysis was circulated to three further researchers (MLJ, RH and AM) for peer validation, feedback and finalisation.

In the Likert scale sub-questionnaire, participants rated the game on such factors as its educational, realism and entertainment value - potential uses of the game that we had in mind when designing it in consultation with stakeholders (Supplementary Material 1). Since this data was only collected for one group (those with real-life farming experience), there was no formal analysis of this data and the data were only plotted and described to qualitatively inform the interpretation of results and evaluation of the game.

## Results

### Study participants

A total of 63 people participated in the study; 32 had not worked in farming or a related field, and 31 had worked in farming or a related field. Of those with farming experience, the majority (30/31) had worked with sheep either as farmers (12/31), stockpeople (8/31), veterinarians (9/31), or in other roles (9/31) such as livestock technicians or in agricultural research or policy (N.B. individual participants often had experience in multiple fields, hence numbers do not total 31). Most of those who shared information about the levels of lameness they had experienced in the flocks with which they had worked said that they had experienced annual lameness levels of between 5 and 10% (13/29).

### Participant recall scores in the game

Participants’ accuracy and recall scores were distributed as expected, permitting deeper analysis of participants’ recall scores (Figure 2). The majority of participants (%) had accuracy scores above 50% (D’Agostino’s test; skew=-1.53, kurtosis = −4.24, p-value = <0.01), indicating that they were not simply ‘cheating’ the game by taking a ‘shotgun’ strategy of marking all or most of the sheep as lame in order to maximise their recall scores. High overall accuracy scores also indicated that our animations of lameness were at least realistic enough for participants to recognise them as lameness, further indicating that variation in recall scores was likely to reflect some level of skill in spotting lameness. Recall scores themselves were normally distributed across the entire percentage range (Figure 2; D’Agostino’s test; skew=-0.12, kurtosis = −0.44, p-value = 0.7), precluding a parametric analysis of the factors influencing participants’ these scores.

**Figure 2:**
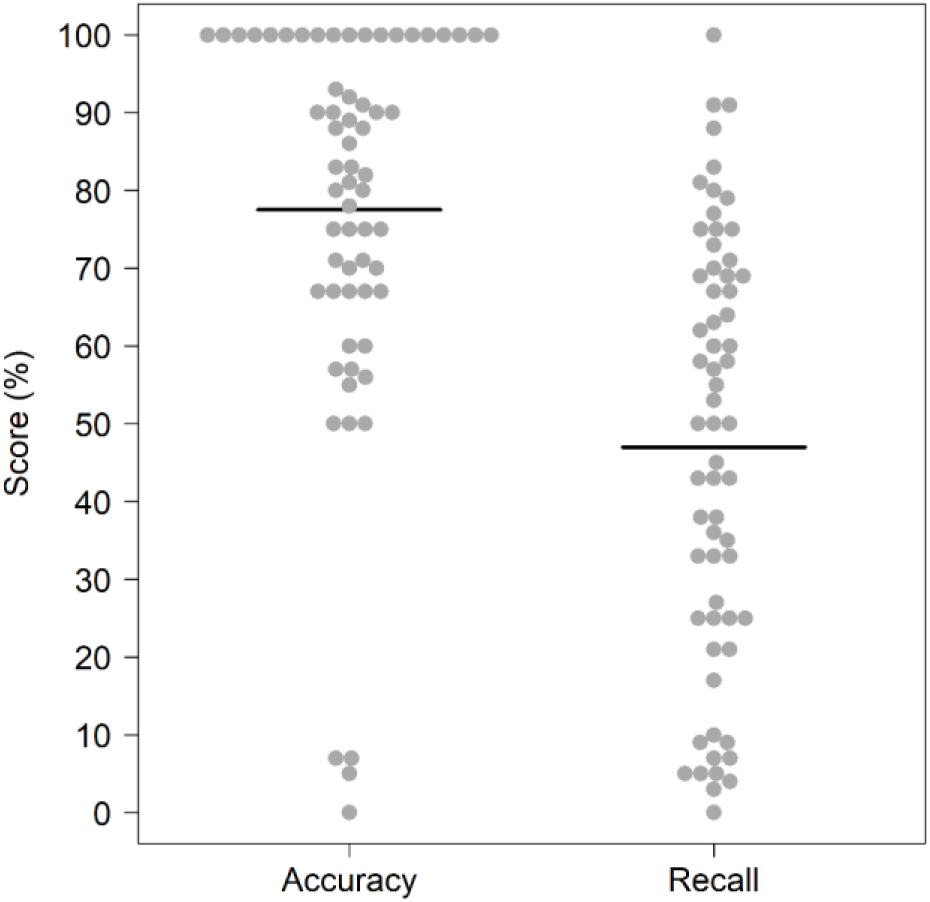
Comparison of distributions of participants’ (n=63) accuracy i.e. number of sheep they marked as lame that were actually lame) and recall (i.e. number of the total lame sheep in the flock that they marked) scores. Individual participant data points are jittered using the beeswarm algorithm (R Package ‘beeswarm’) and mean recall scores are plotted as bold horizontal lines underneath the data points.

#### Farming experience

There was no evidence that real-life farming experience was driving the variation in participants’ recall scores (Figure 3). Initial visual examination of exploratory plots of the data identified no difference in recall scores according to whether participants had ever worked in farming or a related field (Figure 3A). There were also not visually observable relationships between recall and the number of years the participants had spent working with sheep (Figure 3B), or the level of lameness those who had worked with sheep had experienced in the sheep flocks with which they had worked (Figure 3C) - suggesting no higher-level relationships among those with farming experience. Formal statistical testing of the relationship between recall and whether or not the participant had worked in farming or a related field (which encompassed the entire dataset) revealed no significant difference. Those who had not worked in farming or a related field (n = 32) identified a similar percentage of the lame sheep in the game as those who had worked in such fields (n = 31 (‘Farming Experience’ model; R^2^_adj_=0.01, p-value = 0.4, F =0.64, 1 on 61 DF; Supplementary Figure 2).

**Figure 3:**
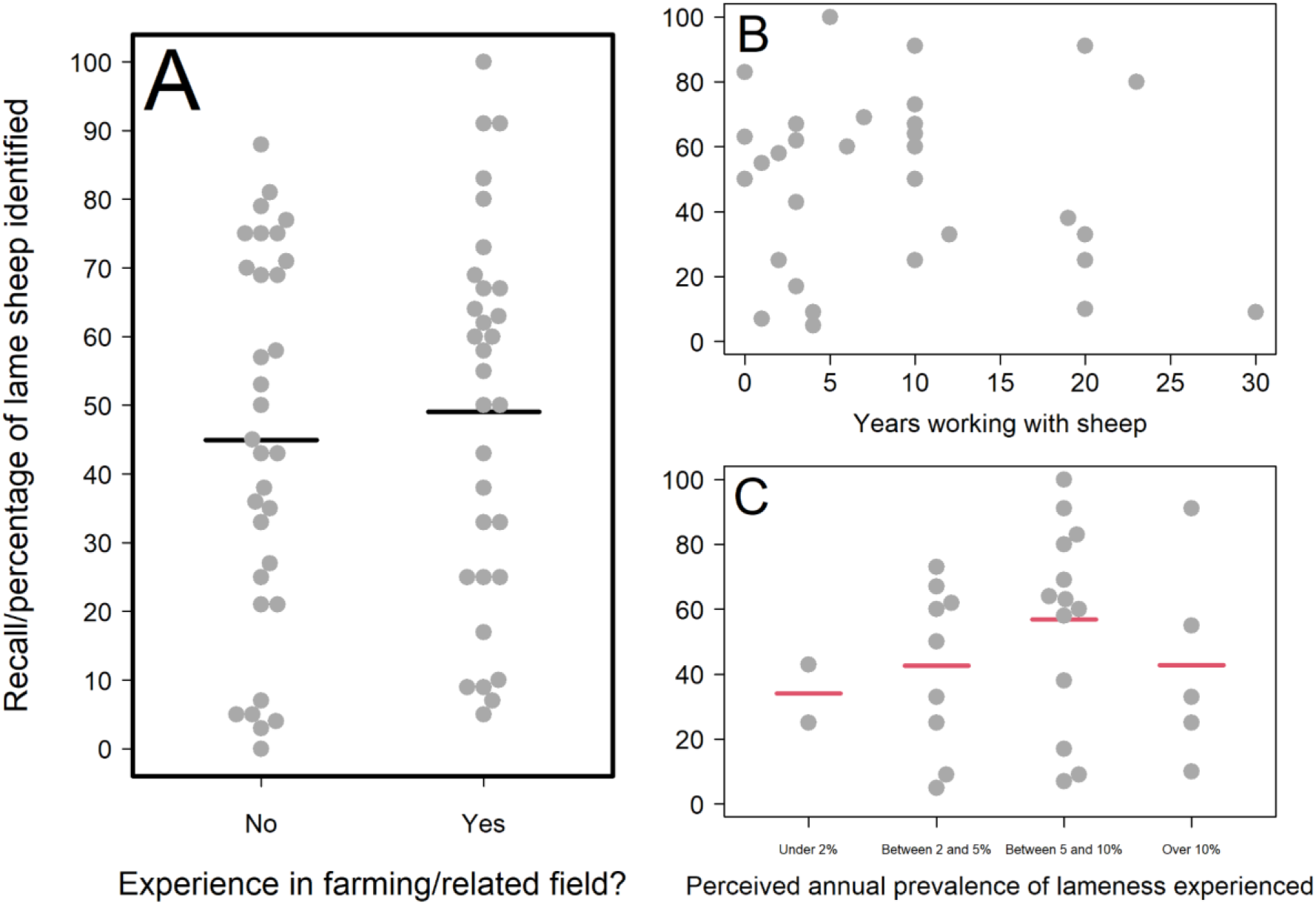
Relationships between participants’ recall scores and their real-life farming experience. A) Recall scores of those without and with farming experience; B) Recall scores and years of farming experience spent working with sheep (for participants with farming experience). C) Recall scores according to the perceived levels of lameness experienced in real-life flocks (for participants with farming experience who answered this question). For categorical variables, individual participant data points are jittered using the beeswarm algorithm (R Package ‘beeswarm’) and mean recall scores are plotted as bold horizontal lines underneath the data points. Mean recall scores coloured red are those likely to be poor estimates due to small sample sizes i.e. the lower or upper quartile exceeds the 95% confidence limits of the mean. The plot is framed in a bold outline if that relationship was formally tested statistically.

#### Lameness signs looked for

The lameness signs that participants looked for when playing the game were not differentiated by whether they had real-life farming experience (Supplementary Figure 3). Those with farming experience tended to more often look for lameness signs, but there was no statistical difference in the distribution of the signs they looked for compared to those without farming experience (X^2 = 4.77 df = 8, p = 0.8). This suggested that there was potential for an effect of lameness signs looked for that was not already captured in the ‘Farming Experience’ model.

However, when we explored this possibility using exploratory data analysis, no such effects were apparent. All of the relationships between in-game recall scores and the signs participants looked for were weak according to initial visual observation of the plotted data (Figure 4). Lameness signs that we included in the animation and deemed to be the most obvious signs of lameness in the game (uneven posture and nodding of the head) were not strongly related to participants’ recall scores. For several signs, the number of participants looking or not looking for the sign was too small to accurately compare the two mean recall scores (red-coloured mean lines). The three relationships with the strongest visual differences in the means were that participants who looked for uneven posture or differing leg speeds (i.e. a limp which we included in the animation as a more subtle lameness sign) scored higher, whilst those who looked for sheep unable to bear weight on a leg whilst standing (a sign of more advanced lameness that was not included in our animation of early lameness) scored lower. However, when statistically tested, neither looking for uneven posture (‘Lameness signs looked for’ model A; R^2^_adj_=0.02, p-value = 0.5, F =1.32, 1 on 61 DF; Supplementary Figure 4), looking for a limp (‘Lameness signs looked for’ model A; R^2^_adj_=0.02, p-value = 0.7, F =1.52, 1 on 61 DF; Supplementary Figure 5) or looking for a raised leg (‘Lameness signs looked for’ model B; R^2^_adj_=0.04, p-value = 0.5, F =2.57, 1 on 61 DF; Supplementary Figure 6) were predictive of participants’ recall scores.

**Figure 4:**
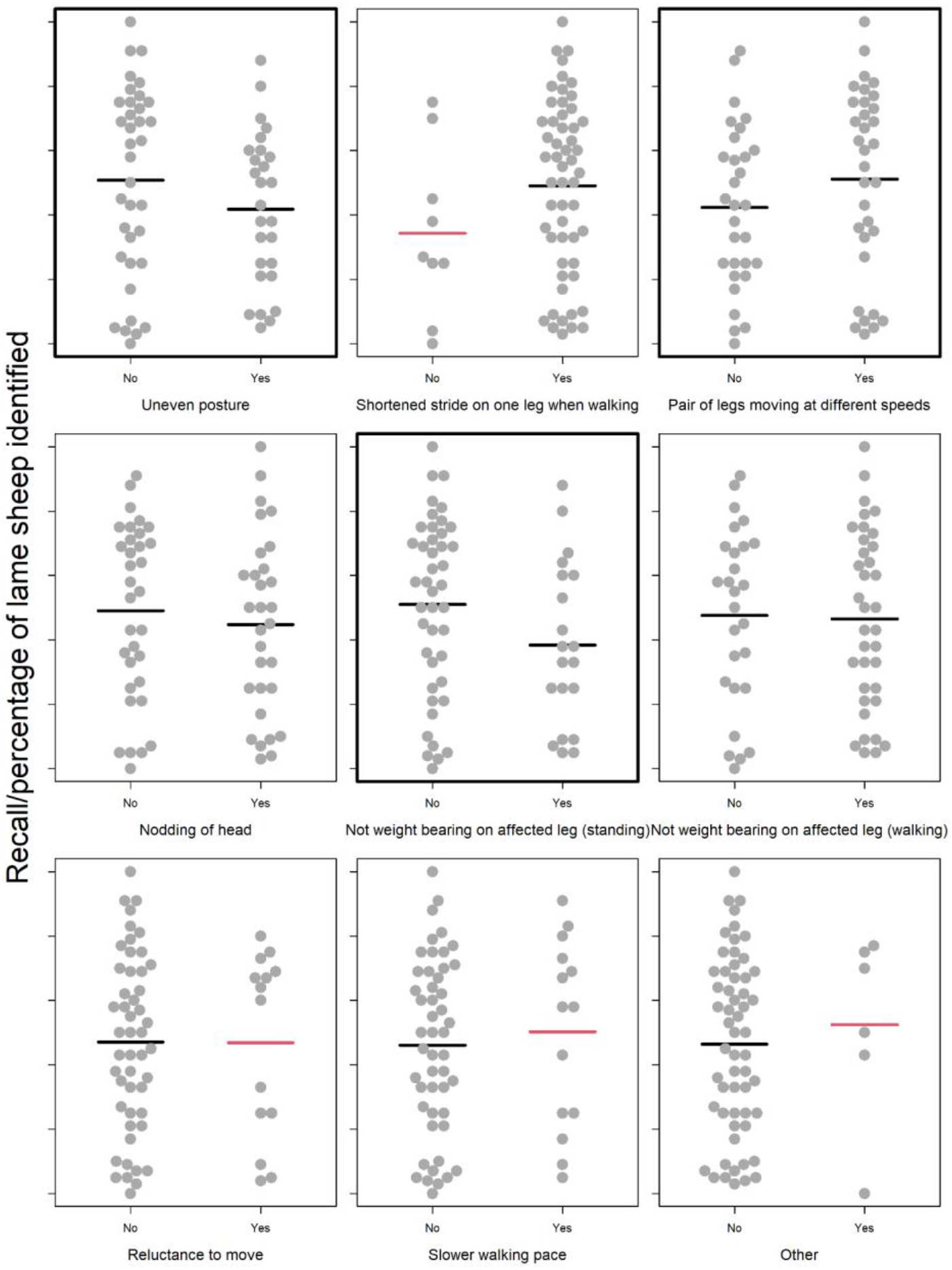
Relationship between participants’ recall scores and the signs they looked for when playing the simulation game. Recall scores of participants that did not and did look for each of 8 classic signs of various stages of lameness (Kaler and Green 2008), plus an extra category of ‘Other’ signs looked for which we asked participants to elaborate upon. For categorical variables, individual participant data points are jittered using the beeswarm algorithm (R Package ‘beeswarm’) and mean recall scores are plotted as bold horizontal lines underneath the data points. Mean recall scores coloured red are those likely to be poor estimates due to small sample sizes i.e. the lower or upper quartile exceeds the 95% confidence limits of the mean. The plot is framed in a bold outline if that relationship was formally tested statistically.

#### User engagement

Similarly to the ‘Farming experience’ and Lameness signs looked for’ variables considered, most aspects of participants’ user engagement did not have a strong effect on recall scores, with recall scores either widely distributed within, or thinly spread across, the explanatory categories considered (Figure 5A-F). The exception to this was that the time spent playing was positively and linearly related to in-game recall (Figure 5G), which formal statistical testing confirmed (‘User Engagement’ model; R^2^_adj_=0.17, p-value = <0.01, F =12.65, 1 on 61 DF; Supplementary Figure 7). Specifically, within the range playing lengths observed (1.45-10 minutes), participants identified an average of 2 additional sheep for every additional minute played.

**Figure 5:**
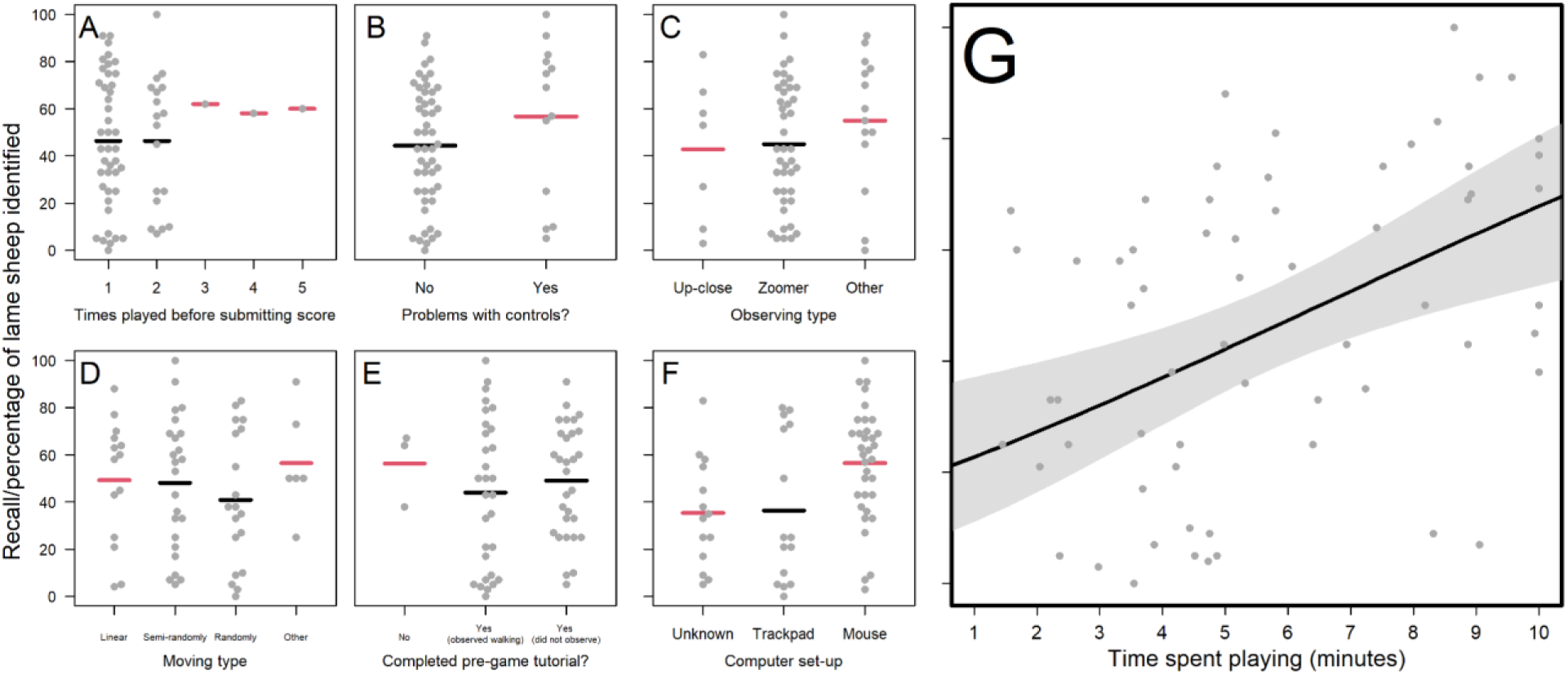
Recall scores of participants according to B) How many times participant played the game before submitting scores; B) Whether participant had problems using the game controls; C) Whether participant observed the sheep up-close, from afar and then zooming in, or using another strategy (e.g. combination of the two); D) How the participant navigated the virtual field to identify sheep; E) Whether the participant completed the pre-game tutorial; F) The participant’s computer set-up/pointing device; G) The time the participant spent playing the game. For categorical variables, individual participant data points are jittered using the beeswarm algorithm (R Package ‘beeswarm’) and mean recall scores are plotted as bold horizontal lines underneath the data points. Mean recall scores coloured red are those likely to be poor estimates due to small sample sizes i.e. the lower or upper quartile exceeds the 95% confidence limits of the mean. The plot is framed in a bold outline if that relationship was formally tested statistically. For a more detailed explanation of what the categories mean (particular for ‘Observing type’ and ‘Moving type’) please refer to Supplementary Material 2

### Feedback on the game from those with real-life farming experience

#### Feedback received as open-form responses

19 out of 31 participants with real-life farming experience provided additional free-text feedback (alongside the Likert scale feedback) on the game and their experience playing it. During the qualitative thematic analysis (Braun and Clarke 2006) of these responses, five key themes emerged: the perceived realism of the game, reflective experiences, challenges of playing the simulation game, emotional responses to the game, and participants’ suggestions for improvement.

##### Perceived realism of the game

Participants with real-life farming experience commented on their perceptions of how realistic the game was as a simulation of real-life experiences with sheep on the farm. Opinion regarding the realism of the simulation was split, with some participants considering that the simulation was *“really realistic”* and *“mimicked sheep well”*, and others expressing that they thought our animations were not sufficiently realistic to enable them to apply their real-life experience of spotting lameness in the game. For example, one participant simply remarked that the simulation was “*not realistic”*, while another noted in particular that “*the main issue was the unrealistic movement of the feet on the ground while standing”* - an animation bug that was known to researchers, but considered minor and impractical to fix before study initiation given timeframe/budget available.

##### Technical challenges playing the simulation game

Participants with real-life farming experience commented on a range of technical challenges relating to the game simulation that hindered their ability to engage with and benefit from the game. Four different aspects were identified as sub-themes: lack of movement of the sheep; simple, unnatural and confusing game simulation of sheep behaviour; inability to mark non-lame sheep; usability and animation/simulation issues.

The first sub-theme, the lack of movement of the sheep, concerned the perceived staticness of the digital sheep and the inability of the player to affect it. Additionally, we considered that the challenge of spotting very subtle signs of lameness efficiently when only presented with glimpses of the behaviour was a key skill to early identification of lameness in the flock. However, as one participant observed, “*lameness is not often identified when animals are static in the field, more often when animals are being moved or handled”*. A key issue for participants appeared to be that that we did not fully simulate the real-life behaviour of farmers *“working the flock”*, whereby the farmer or stockperson moves around and through to flock to stimulate sheep movement: *“I think most farmers would say that they also assess lameness by making the sheep walk / move away from them rather than just wait until they walk*”.

The second sub-theme, the ‘simple, unnatural, and confusing game simulation’, concerned distractions brought about by the games’ computational performance as a consequence of the perceived realism of the game previously described. Commenting on the ‘foot slide’ bug, one participant noted that while “*the sheep animations are good, but to a trained eye I found them confusing, e.g. none of them stood grazing in a normal posture because they were all jiggling their legs all the time”*. In addition to the ‘foot slide’ bug, there were other technical challenges such as game lag and stilted movement, reflecting limitations of the technical systems involved in presenting the game to players online. For example, one participant commented that it was *“sometimes difficult to tell if a normal movement of sheep was a game lag”*, while another considered that the *“movement [was] stilted which made identifying slightly lame sheep virtually impossible*”.

The third sub-theme was the inability to mark non-lame sheep. The fact that there was no means to mark non-lame sheep in the game made it more difficult for participants to remember which sheep they had already assessed, though this was also an intentional design choice. We omitted this feature after discussion with our advisory board, because we considered that in real-life situations of assessing lameness, only lame sheep are usually marked. One participant’s comment composed this theme, mirroring the difficult compromise between playability and realism that we encountered when designing the game: “*It was a bit frustrating not to be able to mark non-lame sheep when surveying, but that is more realistic and requires strategy*”.

The last sub-theme concerned usability and animation/simulation issues. A lack of smoothness in game animations was commented on in particular by one participant who noted that this issue made “*the distinction between a normal walking gait and a limp less easy to discern”*. Meanwhile, another participant noted a lack of clarity in the graphics, which meant *“it was hard to see if they were holding a leg slightly up”*. Another participant also mentioned the ‘foot slide’ bug, which was commonly commented on by participants from a range of perspectives, as reflected in the previous sub-themes.

##### Emotional responses to the game

Participants with real-life farming experience frequently used the open-form feedback request to express how they felt playing the game, with the 5 key sub-themes emerging in thematic analysis: enjoyment, interest, boredom, frustration, and lack of appeal.

Some participants express positive feelings about the game such as enjoyment (sub-theme 1), saying that they *“enjoyed the game”* and found it *“entertaining”*. Others expressed interest in the game (sub-theme 2), with one commenting that it was *“interesting to be looking for signs in virtual sheep”* and another that they “*thought this was brilliant*”.

However, some participants also expressed negative feelings toward playing the game. Boredom (sub-theme 3) and frustration (sub-theme 4) were expressed, and appeared to be mostly related to the staticness of the sheep and their inability to affect it (theme 3: sub-theme 1). For example, one participant noted that they became “*bored waiting for the sheep to move*”, and similarly others commented that the game was *“frustrating”* or *“very frustrating”* to play (sub-theme 4), with one noting explicitly that the cause of their frustration was *“waiting for the sheep to move”*. In addition, one participant expressed a more general lack of appeal (sub-theme 5), such as *“This sort of game doesn’t appeal to me I’m afraid. I’ve always worked in the real world”*.

##### Reflective experiences

Participants with real-life farming experience also reflected on the experience of playing the game and the strategies they employed to identify lame sheep. For example, one participant emphasised how the game *“allowed me to get a better sense of my knowledge and skills”*, reinforcing how the game could enable participants to take stock of their current stockpersonship skills, and serve as a useful benchmarking exercise. However, others found the game too easy as one participant commented that *“lame sheep aren’t always that easy to spot in a field”*, while another commented that “I *think most sheep farmers know the signs of lameness*”. Considering strategies, participants mentioned that in real life, it was important to *“walk around the flock”*, and noted that the sheep *“would move”* in response to the farmer’s movements in a more realistic setting.

##### Participants’ suggestions for improvement

Participants with real-life farming experience also offered suggestions for improvement to the game or to inform future games in this field. These suggestions fell into two broad categories.

Firstly, in line with other feedback, there were suggestions relating to making sheep move, e.g. using additional mechanisms and characters. Creating more natural movement patterns, rather than just a realistic gait, was considered an important priority for future improvement. Participants offered a range of perspectives on how to make the sheep move, but a common view was that it was important to be able to actively move the sheep, as a farmer would in a real-life field, rather than passively waiting for the sheep to move in order to be able to assess gait, as in the current game. For example, one participant suggested: *“If there was a way to make each sheep move, that would really help to keep engagement”*. Meanwhile, another participant suggested adding a sheep dog character to “*run round”* the sheep, while another suggested “*walking a person around so they [the sheep] walk away from you”*. It was commonly agreed that active flock management would be needed for the game experience to be realistic.

Secondly, other participants suggested providing additional visual or sound feedback in the game. One participant commented that visual feedback could be reinforced by offering a *“slightly more realistic depiction of sheep movement for non-lame sheep”*, while another participant considered that auditory feedback regarding the correct identification of a lame sheep, *“maybe a sound…as you chose the correct animals”*, could be a useful addition.

#### Feedback received via the Likert-Scale Questionnaire

Feedback from a Likert-Scale questionnaire suggested that the 31 participants with real-life farming experience could see the potential of games like ours as professional training-type tools in agriculture, but were unsure whether our prototype had realised this potential fully (Figure 6). The majority of participants agreed with statements related to the purpose (“It is clear to me how the contents of the game are related to my profession”; 87%) and usability (“The game rules are easy to understand”; 87%) of the game. Similarly, statements expressing the educational potential of the game - “Learning to play this game was easy” (77%), “The contents and structure helped me to become confident that I would learn with this game” (71%), and “I would recommend this game as a form of training/educational tool” (65%) - received agreement from the majority of participants. However, there was lower agreement with the statement expressing that this educational potential had been achieved (“I feel satisfied with the things that I learned from the game”; 45%). Regarding statements related to the realism of the game, there moderate agreement (56%) with the statement “I achieved the goals of the game by applying knowledge” and low agreement (38%) with the statement “The game is a realistic representation of recognising sheep lameness in the field”. Statements related to the entertainment value of the game received varied responses. Most participants felt the game offered an appropriate level of challenge (68%), and expressed that they had some fun playing (61%). However, many participants appeared to find the game boring by the end of playing; expressing that they felt the game became monotonous as it progressed (55%) and not recommending it as a form of entertainment (48%). The game was not deemed particularly absorbing, as reflected by the fact that most participants did not lose track of time (77%) or forget about their immediate surroundings (58%) while playing the game.

**Figure 6:**
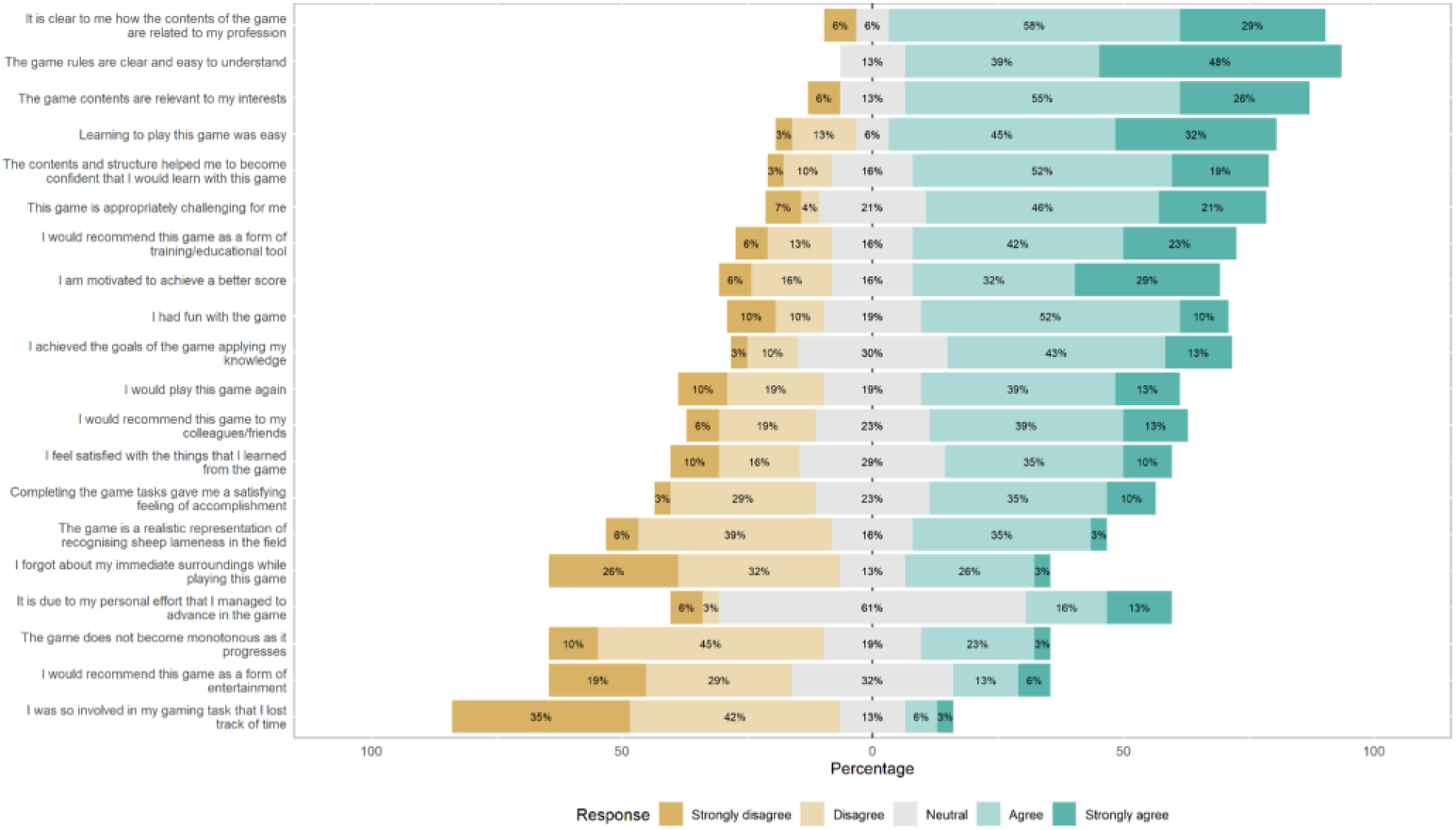
Quantitative feedback given on the game via a Likert Scale questionnaire. Statement rated are shown on the rows, with the total percentages of participants with farming experience responding negatively, neutrally and positively to the statements overlayed on the stacked bar graph.

## Discussion

Our online evaluation study highlighted the challenges and opportunities of using simulation games for the purposes of supporting real-life livestock husbandry practices. Whilst positive feedback from participants indicated signs of potential for using simulation video games in this context, barriers to this audiences’ user engagement with computer games like ours hindered this potential from manifesting more widely. Particular barriers included participants’ apparent desire for high levels of realism and engagingness in the game - expectations which we struggled to meet and therefore limited the game’s ability to function as a tool for quantitatively assessing, train and understand farmers’ ability to recognise the earliest, subtlest signs of lameness. Nonetheless, the results of the study provide valuable insights for the design and use of future similar games and studies in livestock husbandry.

### User engagement shapes in-game performance where participants struggle to relate to the simulated environment

There was substantial variation in participants’ recall scores that was not well explained by the metadata about participants that we collected via the after-game questionnaire. Thus, even if effects of real life farming experience or the lameness signs looked for were influencing participants’ recall scores, they may have been outweighed by the effects of these unknown causes of variation. Whilst it is hard to guess what these causes are, the finding that the time spent playing was the only driver of participants’ in-game performance, alongside the results of our qualitative analyses, suggest that the results were at least partly due to participants not finding the game sufficiently realistic or engaging.

Regarding realism, although participants’ explicit statements about the game’s realism were split, many of the other themes identified in participants’ feedback related back, in some way, to the game not sufficiently reflecting real life. Statements expressing the realism of the game were also generally disagreed with in the Likert-Scale questionnaire, suggesting most participants had some issues with the realism of the simulated experience. Our pursuit of realism during the game development process was heavily motivated by early interviews with farmers, who were the intended audience of the game (Supplementary Material 1). Although our sample size of potential users was small and may not be reflective of all the potential users of such games, there was a consistent feeling among interviewees that a research/education game of this sort should reflect real-life scenarios as accurately as possible. However, the difficulties we faced in achieving this desired level of ‘realism’ probably limited the game’s potential as tool for training or assessing farmers’ lameness recognition skills direct. Certainly, some level of realism was achieved; the high accuracy scores of all participants indicated that participants could recognise our virtual lame sheep as lame (Figure 2). However, the lack of an expected difference in recall scores between those with and without farming experience, alongside the lack of an effect of the lameness signs looked for, suggests that our animations were perhaps too obvious. As one farming-experienced participant’s feedback attested to, in the field sheep behaviour is much more complex (e.g. hiding weaknesses from farmers as part of their prey instinct), and farmers look for a wide variety of body language cues when they observe a sheep’s gait for lameness beyond just the textbook examples.

Another possible reason recall scores in the game failed to reflect real-life experience and skills is that the game was not sufficiently engaging for participants to play. Some participants expressed boredom or frustration in the after-game feedback, which is probably the reason many quit the game early (reflected by the wide range of times spent playing in Figure 5). Again, this was partly related to realism; in the pursuit of realism, we probably made the game overly long and sacrificed entertainment value. For example, the decision to program the sheep to only walk intermittently to better reflect real life behaviour lead us to develop a game that was 10 minutes long to ensure participants had a sufficient opportunity to observe each of the 25 sheep in the virtual flock walking at least once. Especially considering that the game consisted of repeating one task, this may have caused many participants to quit the game early, impacting their recall scores. Although an overemphasis on the “fun factor” can be detrimental to the use of games in non-gaming contexts like agriculture (Monk 2002), game-based approaches must still achieve a user experience that is to some extent playful and engaging (Treiblmaier, Putz, and Lowry 2018), especially as many people hold preconceived notions that video games are always designed for the purpose of entertainment. More technical problems such as in-game ‘bugs’ and problems participants had engaging with the virtual flock may have further limited the game’s engagingness. Again reflecting of the minutiae of signals that farmers process when trying to recognise lameness, in-game malfunctions such as the foot-sliding ‘bug’ - which we considered relatively inconsequential and not a priority (in terms of what was feasible given the predetermined project budget and time frame) to fix before the study roll-out - turned out to be quite distracting for some participants. More generally, the inability to move the virtual sheep and ‘work the flock’ was frustrating for some participants, who expressed that passive observation was not an efficient way to identify lameness.

Finally, we would like to highlight the importance of budgetary limitations in limiting our ability to achieve the levels of realism and engagingness that participants expected. Although we worked with a skilled game programmer and animator experienced in scientific animation, we were not always able to make the most of their skills due to the constraints of our £5000 budget (Supplementary Material 4). This limited the time the game programmer and animator had available to work on the project, and they were thus not always able to make use of the feedback and support that was available from the review and testing stages (e.g. addressing boredom issues or the ‘foot slide’ bug). Furthermore, funding was not sufficient to enable us to hire someone with subject-specific expertise (e.g. a sheep farmer) to directly work with the game developer and animator on a day-to-day basis (which they expressed would have helped). We therefore strongly recommend that future grant applications for serious game projects seek sufficient funding to cover more of the primary game developers’ time and also facilitate much closer, more direct collaboration between the game developers and the game’s intended audience. This would enable design choices to be driven by the intended audience’s involvement and not by what is feasible due to budget limitations, increasing game acceptance and the potential benefits of this medium.

### Insights on lameness recognition practices

Our study did reveal some interesting insights on lameness recognition and produce some evidence of future potential for using games as a tool in livestock husbandry education and research.

Firstly, our inter-disciplinary study points to the way in which animal ailments like lameness may resist precise scientific definitions. Despite the highly controlled *in silico* laboratory we created in which lameness is precisely programmed into the virtual flock, we nonetheless observed a wide variety of recall scores. Although we primarily attribute this to the effect of time spent playing (supported by our quantitative analysis) and the difficulty of adequately mimicking real-life in a video game (supported by our qualitative analysis), our results are also likely to reflect the inherent subjectivity involved in assessing lameness. Previous research has shown that even when observing (videos of) real sheep, farmers and other specialists vary substantially in what they define as lame (especially for early lameness), with different ‘thresholds’ for defining lameness and acting upon it (Kaler and Green 2008). Thus whilst *“most sheep farmers know the signs of lameness”*, as one participant commented, lameness is a spectrum that may resist a precise definition and be tied up with individual farmers’ lived experience. The use of mixed methods reveals this acutely, lending a unique level of support to the hypothesis that subjective experience must be better considered when seeking to design interventions for livestock husbandry issues like lameness in farming.

Similarly, some of our results suggest that the game produced a level of understanding that would not have been so easily achieved with solely survey-based methods, allowing farmers to engage with researchers in novel ways. In particular, we note that the process of researchers illustrating (through the creation of a game) their ‘vision’ of what lameness recognition on the farm looks like (and requesting feedback from those with real-life farming experience on this) facilitated conversations about lameness that perhaps may not have happened with solely survey-based methods - one of the main benefits of the human-centered design approach. Participants reacted strongly to the artificial, simplified world we created, telling us what was missing from our vision and highlighting the limitations of our understanding as academics, proving the utility of iterative prototyping (Lim, Stolterman, and Tenenberg 2008). A notable example of this was those with real-life farming experience questioning our assumption that early lameness recognition depended on passive observation and making clear that it depends on actively ‘working the flock’. Similarly, participant feedback and performance data suggesting that the game easy revealed how academics might misdiagnose real-life problems (and by implication, prescribe flawed solutions); revealing that the decision-making challenge in lameness management may not lie in being able to recognise lameness early, but in being able to act upon this knowledge accordingly (e.g. in finding time and resources to catch and treat sheep). Such assumptions may not have been obvious in a less creative, interdisciplinary project, and has implications for managing lameness in the real-world; suggesting that finding ways to embed lameness reflection and monitoring into existing shepherding practices might help reduce lameness more than trying teach farmers the signs of lameness.

Finally, on a more fundamental level, the game-based, incentivised study appeared to function well as a ‘hook’ to encourage agriculturalists to discuss and participate in a more conventional survey about managing animal health. Many participants shared positive feedback on the game, especially with regards to its potential as an educational tool (even if this had not been fully realised). Furthermore, anecdotally at least, some agriculturalists suggested that the novelty of using a game made the study more appealing (especially when compared to solely survey-based studies that they often get requests to participate in). The game also supported experiential learning through reflection and facilitated the acquisition of up-to-date information on lameness recognition in UK farmers. Agriculturalists were clearly at least trying to spot lameness in the virtual sheep as they would for real-life sheep, and some explicitly expressed that it allowed them to take stock of their real-life practice. The fact that those with farming experience tended to look for lameness signs more often (Supplementary Figure 3) is consistent with the previously reported finding that most farmers know how to identify lameness (Kaler and Green 2008), though a larger sample size would be needed to confirm this. New sociological tools like games may therefore at least help facilitate survey methods and encourage more active participation and engagement between farmers and researchers, as well as support learning through reflection.

### Implications for use of games in livestock husbandry

Our findings have important implications for the future development of games intended as tools to engage with farmers on livestock husbandry issues such as lameness and stockpersonship. In particular, they highlight that future similar projects should consider carefully whether games are best used as ‘virtual laboratories’ to study and train participants, or more as tools to facilitate discussion between researchers and stakeholders in livestock husbandry.

If the games being developed and/or used are intended to be used as ‘virtual laboratories’, researchers should consider carefully whether the levels of both realism and engagingness that we expect farming audiences to demand of this medium are achievable before initiating the project. A bigger budget, resources, experience and closer engagement with farmers, stockpeople and farm vets will certainly help to tap the full potential of games in this context - though balancing realism and engagement is still likely to be a challenge when using this medium with this audience. One approach that might prove a particularly fruitful avenue for exploration in this regard is to build on existing games, rather than creating games anew. This was the ethos of a recent study that demonstrated the educational value of games in learning natural history by using the professionally-developed video game Red Dead Redemption 2, leveraging its established popularity, realism and entertainment value to engage participants whilst saving time and resources (Crowley, Silk, and Crowley 2021). The hyper-real popular video game Farming Simulator - which is already played by farmers (Lane 2018) - might serve a similar role in future studies of games in agriculture. Indeed, Pavlenko et al. (2021) have already had some success building a ‘mod’ (a ‘modification’ - new game content/software created by someone other than the primary game development team) for this game to encourage the adoption of precision agriculture technologies. Alternatively, future projects might do better to use real-life imagery rather than 2/3D models to simulate agricultural environments; this ethos is already being successfully deployed by the ‘3D farms’ project centered around virtual reality to overcome logistically and accessibility challenges in agricultural training (Barber 2016).

If games are intended to be used more broadly as tools to facilitate discussion between researchers and stakeholders, researchers should be less tied to realism and be more open to letting the game develop organically in close consultation with stakeholders. The game development process itself may facilitate knowledge exchange more than end product, as evidenced by the insights on real-life lameness recognition practices gained through participants telling us what our game was missing, for example. This is something that should be explored in a more dedicated way in future studies using games to engage with farmers on aspects of livestock husbandry.

## Conclusions

The use of games in agricultural research has been increasing in recent years and here, we attempted to develop and use a game to support the study of lameness recognition in UK sheep farmers. We found that besides the positive effects of the game in supporting understanding, knowledge exchange and reflection of lameness, difficulties engaging the agricultural audience limited the potential of the game for education and research. In particular, experienced livestock farmers, stockpeople and veterinarians requested much higher levels of realism and engagingness than could be achieved with the limited project budget and time-frame.

These results suggest that more needs to be done to establish whether games can be a cost-effective tool in livestock health education and research, and to explore the most effective ways and scenarios in which to use them. Future similar studies should seek to obtain larger budgets, build on existing agricultural simulation games, and work more directly with their target audience, in order to develop games that can more acutely address the challenges of managing livestock health in the twenty-first century.

## Acknowledgements

We would like to thank to all the farmers, veterinarians, academics and others who participated in the human-centered design, piloting and evaluation process to create these game and execute this study. Thank you to our advisory board who helped shape the game, animations, and study design. Thank you to the Health and Environment Public Engagement Group at University of Exeter’s European Centre for Environment and Human Health, who acted as test participants before the formal roll-out of the online study. Thank you to ethics and press office staff at the University of Exeter for enabling the study to meet ethical guidelines and advertising the study in order to recruit participants. Finally, many thanks to the 63 participants who gave their time to take part in the online evaluation study, whose data forms the basis of this paper. The last author would also like to acknowledge the Centre for Artificial Intelligence, Robotics and Human-Machine Systems (IROHMS) operation C82092, part-funded by the European Regional Development Fund (ERDF) through the Welsh Government. This study was funded by the GW4 Crucible Seed Funding Award Cru20_3.

## Data availability statement

The anonymous data collected during the online evaluation study (the results of which are reported in this paper) are freely available in raw (direct output from MS Forms platform) and formatted (tidied using R code) formats at [INSERT OPEN SCIENCE FRAMEWORK LINK UPON ACCEPTANCE OF MS] and the lead author’s Github repository (https://github.com/befriendabacterium/lamenessgame). R code to reproduce the analysis and figures reported in the manuscript is also deposited here. The Supplementary Materials of the manuscript also contain the thematic analysis of the open-form feedback in a more accessible Word document format. All participants approved the open publication of these anonymous data when signing the consent form to participate in the study.

## Supplementary Material

### Supplementary Material 1: Development of the Game

The game was developed using a human-centered design process that began in June 2020, when our interdisciplinary group of researchers (with expertise in microbiology, engineering, social science, and human-computer interaction) started a small project (with a budget of £5000) to initially explore the potential use of game-based approaches in the context of antibiotic use practices in livestock production (M. Jones et al. 2020). There were three main phases of development.

#### Phase 1: Gathering initial requirements and responses to early ideas about a game about antibiotic use in livestock production

During the initial requirement gathering stage, we conducted interviews with 3 farmers (2 cattle/sheep, 1 pig) and 1 farm vet recruited through the JustFarmers platform (A. Jones 2022). Interviews were ethically approved by Cardiff University School of Informatics Ethics Committee, and comprised two parts - a first part of the interview attempting to understand the participant’s experiences and challenges managing disease in livestock production including the current use of antibiotic in livestock production, and a second part in which we started to explore the design space using iterative prototyping to visualise and communicate two early prototypes (Supplementary Figure 1A) to explore their possibilities, and limitations (Lim, Stolterman, and Tenenberg 2008), as well as to provoke discussions and look for alternative ideas (Fallman 2008). The first design exploration was a wireframe of a simple game intended to communicate the balance between disease prevention and antibiotic stewardship on the farm (prototype 1a). The second design exploration was a slightly higher fidelity wireframe of a game in which players have to judge which animals to treat with antibiotics by selecting animals and assessing their list of symptoms provided as a journal entry (prototype 1b). These wireframes were demonstrated to the 4 interviewees in online meetings, in order to broadly introduce the concept of a game centered around appropriately treating sick animals with antibiotics; to stimulate discussion with farmers about whether such games might be useful for education (farmers were shown the wireframes via screenshare and asked for their thoughts); and to act as conversation-starters. Participants discussed the early prototypes and provided their feedback on their utility.

The interviews and feedback sessions around the prototypes with farmers and vets were recorded and thematically analyzed to identify major themes and ideas to inform the game’s development (Braun and Clarke 2006). One of the major findings was that farmers felt that in seeking to make a game to support the reduction of antibiotic use in livestock farming, an explicit emphasis on antibiotic use practices was not necessary. Interviews consistently indicated that antibiotic use practices in livestock production are underpinned by stockpersonship in animal health management and the farmer’s challenges and ability to early recognize animal behavioral signs and physical characteristics of sick animals. Regarding the first version of the prototype, participants highlighted the importance of the realism of the game in relation to the natural surroundings of the farm and the animals. In addition, participants highlighted how unrealistic visual elements (cartoonish looking; Supplementary Figure 1) of the prototype can be distracting. Overall, participants suggested that a fruitful avenue to pursue would be to develop game with a realistic-feel that served as a stockpersonship training tool, expressing the sentiment that being able to spot disease early was more critical and challenging than knowing how to treat it.

#### Phase 2: Development of first playable prototype (Where’s Woolly?)

Building on the findings of the first phase, in the second phase we focused on developing a more higher-fidelity prototype game focusing on stockpersonship within a sheep farming context (given that the stakeholders we were engaging had shared experience working with sheep). The game was loosely intended to support antibiotic stewardship in agriculture by providing an environment for testing, honing and studying farmers’ ability to recognise the early signs of ill health in their livestock - though we remained open to other potential uses of the game throughout our evaluation process.

The prototype we developed (prototype 2) was a game in which players were presented with three scenarios of identifying sick sheep in a flock e.g. identifying animals that were walking slower than other animals, or standing apart from the flock, or not eating (Supplementary Figure 1B). Scenarios were helpful to illustrate the potential and future use of the game as well as to gather feedback and identify potential problems (Bødker 2000). To add more realism to the game, this prototype was created using the Unity development platform (Unity Technologies 2021) that facilitated the creation of a 3D virtual environment containing more details such as a grassy terrain, bushes, trees, and more realistic models of the animals (in this case we used an existing sheep model (‘asset’) considering the previous feedback). Seven participants including a sheep/cattle farmer, a veterinary microbiologist, 2 healthcare academics from our networks and 3 of our own team members were asked to play and provide feedback via a Likert Scale questionnaire to explore initial playability of the game, adapted from the MEEGA+ method (Petri, Gresse von Wangenheim, and Borgatto 2017), and a short usability questionnaire focussed on identifying in-game bugs and gathering technical suggestions for improvements. Healthcare academics from our networks and our own team members obviously could not offer a perspective on the game based on real-life farming experience and were more subject to bias in their evaluation, which probably limited this evaluation. However, given the goal of understanding the potential playability of the game and identifying technical issues and fixes, this was less of a concern at this stage.

No formal qualitative analysis was conducted on these data due to the brevity of the information provided. Briefly though, participants provided positive comments about the game’s potential for training and research but also highlighted the need for the game to be more realistic. For example, participants suggested that we seek to include in the game more sheep showing subtle symptoms and provide feedback on whether the animal was treated correctly. Participants also suggested that we include an introduction screen to explain the different roles and game actions to the players as well as different camera angles. Overall, participants recognised the game’s potential to improve livestock health management, especially as an educational tool for inexperienced farmers.

#### Phase 3: Development of final prototype (The Lameness game)

Building on the results of Phases 1 and 2, we chose to develop a game focused on lameness recognition in sheep farmers (Supplementary Figure 1C). Lameness was chosen as the theme of the final prototype not only because it provided a focal point for developing a more realistic game, but because it was closely intertwined with stockpersonship, resonated with many of the (mainly sheep and cattle) farmers and vets we consulted, and is a key challenge in UK livestock farming with wide-ranging implications for productivity, welfare and antibiotic stewardship.

For the development of the final prototype that was evaluated in this study, we first enrolled an animator (TL) with experience with scientific animation to work with our game programmer (OM), focusing on developing a realistic animation of lame and non-lame sheep which could form the basis of a game to test farmers’ lameness recognition skills. This was done through a mix of consulting scientific source materials, written and video, mainly from Kaler and Green (2008), scientific experts and producing our own reference material (co-author HV filming her own sheep). This information was used to modify an existing 3D sheep model and its animations purchased from the Unity Assets store (Red Deer 2020), which was then integrated into the game. We created an expert advisory panel of farmers and sheep lameness academics, including some of the co-authors. The first author conducted a one-hour focus group to consult with stakeholders and receive feedback on the animation, aesthetics, gameplay mechanisms and future refinements. Notes were taken during the consultation sessions with stakeholders, which informed the development of the game (though no formal thematic analysis was conducted due to time and resource constraints). Feedback from the advisory panel emphasized the need to improve the sheep gait animations, which we responded to by investing more time and resources into animation refinement and their smooth integration into the game.

### Supplementary Material 2: Questionnaire

#### Consent

1. By checking this box I confirm that I have understood and agree with all of the above statements and I consent to taking part in this project. You must tick this box to agree with all of the above statements, in order to part in the questionnaire.

#### Game Results

You will need to record your game time & scores after playing the game so please read the instructions below carefully before playing:

STUDY INSTRUCTIONS:

1. Go to https://wheres-woolly.itch.io/lameness-game, leaving this form open
2. Play the tutorial, the afterwards play the Game itself
3. Upon finishing the game DO NOT CLOSE THE WEBPAGE - you will be shown your scores (an example screenshot is shown above) - keep it open and enter your scores in the form below, then continue with the rest of the questionnaire.

REMINDER - GAME RECOMMENDATIONS:

- Desktop or laptop computer - The game should not be played on touchscreen devices (i.e. smartphone or tablet).
- Mouse with a scroll wheel or a laptop trackpad - to ensure efficient game-play.
- We recommend playing the game in one of the following web-browsers: Microsoft Edge, Google Chrome or Mozilla Firefox (other browsers are not supported)
- If the game is running slowly, try closing unused web-browser tabs (not this one)

2. Time remaining on clock when you ended the game by clicking ‘Done (in the format nnn seconds e.g. 596 seconds in the example screenshot).
3. Lame sheep identified (%) - e.g. 0 in the example screenshot
4. accuracy (%) - e.g. 0 in the example screenshot
5. How many times did you play the game before getting these results (0 = it was my first time playing)?
6. Did you play the tutorial before playing the game?

- Yes, and observed the sheep walking
- Yes, but didn’t observe the sheep walking
- No
7. What computer hardware did you use to play the game? (select all those appropriate)

- Laptop or Desktop computer
- Mouse with scroll-wheel
- T rack-pad with pinch zoom
- Smartphone or tablet
- Other
8. Did you experience any problems using the controls or playing the game?

- Yes
- No
9. If yes, please specify

#### Game Strategy

10. What was your strategy for observing the sheep (tick all that apply)? [Data plotted in Figure 5C]

- Observed the whole flock and then zoomed in when I saw one that looked lame [Denoted as ‘Zoomer’ in Figure 5C]
- Observed each sheep up-close until I could see whether or not it had a sign, then moved onto the next sheep [Denoted as ‘Up-close’ in Figure 5C]
- Other (please provide details below) [Denoted as ‘Other’ in Figure 5C]
10. Please provide brief details about your strategy for observing the sheep?
11. How did you move from sheep to sheep (tick one)? [Data plotted in Figure 5D]

- Randomly [Denoted as ‘Randomly’ in Figure 5D]
- Semi-randomly [Denoted as ‘Semi-randomly’ in Figure 5D]
- Started at one end of the flock and worked my way to the other [Denoted as ‘Linear’ in Figure 5D]
- Other (please provide details below) [Denoted as ‘Other’ in Figure 5D]
12. Please provide brief details about how you moved from sheep to sheep
13. What signs did you look for to find the lame sheep (tick all that apply)

- Uneven posture
- Shortened stride on one leg when walking
- Pair of legs which were moving at different speeds
- Nodding of head
- Not weight bearing on affected leg when standing
- Not weight bearing on affected leg when walking
- Reluctance to move
- Slower walking pace
- Other
14. If you answered ‘Other’, please provide brief details about what signs you looked for to find lame sheep

#### Real-world experience

15. Have you ever worked in farming or a related field (e.g. farm vet)?

- Yes
- No
16. How many years have you worked with sheep?
17. In what roles, if any, did you work with sheep (e.g. farmer, stockman/woman/person, veterinarian)?

- Farmer
- Stockman/woman/person
- Veterinarian
- Other
18. If you answered ‘Other’, please provide some brief details about the role(s) in which you worked with sheep
19. What do you think was the average level of lameness in the flock(s) with which you worked/work, over one year?

- Under 2%
- Between 2 and 5%
- Between 5 and 10%
- Over 10%

#### Game Feedback

Please fill in the table below with an indicating how strongly you agree with the preceding statement with 5 being strongly agree and 1 being strongly disagree

How strongly do you agree with the following statements?

1. The game is a realistic representation of recognising sheep lameness in the field
2. Learning to play this game was easy
3. The game rules are clear and easy to understand
4. The contents and structure helped me to become confident that I would learn with this game
5. This game is appropriately challenging for me
6. The game does not become monotonous as it progresses
7. I am motivated to achieve a better score
8. Completing the game tasks gave me a satisfying feeling of accomplishment
9. It is due to my personal effort that I managed to advance in the game
10. I feel satisfied with the things that I learned from the game
11. I would recommend this game to my colleagues/friends
12. I had fun with the game
13. I would play this game again
14. I would recommend this game as a form of entertainment
15. I achieved the goals of the game applying my knowledge
16. I would recommend this game as a form of training/educational tool
17. I was so involved in my gaming task that I lost track of time
18. I forgot about my immediate surroundings while playing this game
19. The game contents are relevant to my interests
20. It is clear to me how the contents of the game are related to my profession

### Supplementary Material 3: Thematic analysis results

#### List of comments for qualitative analysis (note the typing errors from the original, should mark as [cic] if quoting in text)

Participant 8: If there was a way to make each sheep move, that would really help to keep engagement, I got bored waiting for the sheep to move unfortunately.

Participant 15::) [Happy face]

Participant 17: I think most farmers would ay that they also assess lameness by making the sheep walk / move away from them rather than just wait until they walk.

Participant 18: If you wanted to complete the game in a shorter time, you would want the sheep to move around more. I got bored waiting for them to walk. Needs a dog to run round them!

Participant 19: Game animations were not smooth, making the distinction between a normal walking gait and a limp less easy to discern. This scenario may not be very representative, as in my experience lameness is not often identified when animals are static in the field, more often when animals are being moved or handled.

Participant 22: Lame sheep aren’t always that easy to spot in a field

Participant 24: Would be good to get sheep to move, maybe by walking a person around so they walk away from you as it is difficult to assess them systematically.

Participant 27: i got annoyed waiting for the sheep to move. in a flock i would walk around them and the sheep would move.

Participant 34: The graphics werent very clear - it was hard to see if they were holding a leg slightly up. In reality you would move the sheep to look for lameness

Participant 35: I would have enjoyed this game better if the controls worked better the sheep animations are good, but to a trained eye i found them confusing, eg none of them stood grazing in a normal posture because they were all jiggling their legs all the time

Participant 36: This sort of game doesn’t appeal to me I’m afraid. I’ve always worked in the real world.

Participant 38: none [Cannot include in the analysis]

Participant 43: it was entertaiing but i felt there could be improvements made as you chose the right animals maybe a sound so you know your going the right way or a counter in the corner

Participant 44: took a long while for the seep to start moving in the tutorial that i wondered if it was going to move, but I think that’s the point of the questions asking about if I watched the sheep move. i enjoyed the game as it allowed me to get a better sense of my knowledge and skills. it mimicked sheep well but was sometimes difficult to tell if a normal movement of sheep we a game lag.

Participant 50: I thought lameness was really realistic-but was expecting more variation (ie from very early to very severe, different legs, etc - though maybe I didn’t spot that!)

Participant 51: Found it very frustrating. Not realistic. Movement stilted which made identifying slightly lame sheep virtually impossible. Most of the time all sheep standing still, leading to frustration with the game and rushing.

Participant 57: I thought this was brilliant. It was a bit frustrating not to be able to mark non-lame sheep when surveying, but that is more realistic and requires strategy. The main issue was the unrealistic movement of the feet on the ground whilst standing. On my PC there was a foot slide effect. I didn’t look for standing signs as I thought they were more graphics errors

Participant 67: Could be enhanced by slightly more realistic depiction of sheep movement for non-lame sheep

Participant 68: It’s interesting to be looking for sign in virtual sheep, but I got frustrated that I was not able to make them move as would be the case in real life.

Participant 70: very basic. would be nice to have a method of encouraging sheep to move. In real life I would walk around the flock and observe they way the moved. In this game the sheep were fairly stationary which made that hard.

#### First reviewer’s (M.S.B) comments on second reviewer’s (N.V.D) analysis and points of difference

The two analyses presented are compatible to a large extent and reflect far more commonalities than fundamental points of difference. Where there were discrepancies, these reflected different professional backgrounds and differential prioritisation of aspects of the dataset, especially relating to technical versus experiential aspects.

M. S.B. identified 4 themes:

1. Challenges of identifying lameness
2. Psychological responses
3. Realism of farming simulation
4. Technical performance

N. V.D. identified 5 themes

1. Perceived Realism of the Game
2. Reflective experiences
3. Challenges of the Game simulation
4. Emotional Responses to the Game
5. Participant’s suggestions for improvements

I consider that N.V.D has captured the content of my themes, with the following comments.

M.S.B. opted to sort the themes alphabetically. N.V.D. has not stated a logic for ordering the themes. I would prefer to retain alphabetical ordering (unless a strong rationale to the contrary is provided).

I would prefer to retain the theme title ‘Psychological responses’ rather than ‘Emotional responses’, but am happy to add ‘to the game’. I consider that the term ‘Psychological’better captures the range of sub-themes.

I consider the only amendments needed to N.V.D.’s coding are to order alphabetically and to replace ‘emotional’ by ‘psychological’

#### Results/themes identified

1. Perceived realism of the game (PR)

- Quote 1: “the sheep animations are good” (Participant 35)
- Quote 2: “it mimicked sheep well” (Participant 44)
- Quote 3: “I thought lameness was really realistic…” (Participant 50)
- Quote 4: “Not realistic.” (Participant 51)
- Quote 5: “The main issue was the unrealistic movement of the feet on the ground whilst standing” (Participant 57)
2. Technical challenges playing the simulation game (TC)

· Sub-theme 1: Lack of movement of the sheep
○ Quote 1: “: I think most farmers would say that they also assess lameness by making te sheep walk / move away from them rather than just wait until they walk” (Participant 17)
○ Quote 2: “as in my experience lameness is not often identified when animals are static in the field, more often when animals are being moved or handled.” (Participant 19)
○ Quote 3: “I got annoyed waiting for the sheep to move. in a flock i would walk around them and the sheep would move.” (Participant 27)
○ Quote 4: “In reality you would move the sheep to look for lameness” (Participant 34)
○ Quote 5: “took a long while for the sheep to start moving in the tutorial that i wondered if it was going to move, but I think that’s the point of the questions asking about if I watched the sheep move” (Participant 44)
○ Quote 6: “Most of the time all sheep standing still, leading to frustration with the game and rushing” (Participant 51)
○ Quote 7: “I was not able to make them move as would be the case in real life.” (Participant 68)
· Sub-theme 2: Simple, unnatural, and confusing game simulation of sheep’s behaviour (SB)
○ Quote 1: “the sheep animations are good, but to a trained eye i found them confusing, eg none of them stood grazing in a normal posture because they were all jiggling their legs all the time.” (Participant 35)
○ Quote 2: “was sometimes difficult to tell if a normal movement of sheep we a game lag.” (Participant 44)
○ Quote 3: “was expecting more variation (ie from very early to very severe, different legs, etc - though maybe I didn’t spot that!)” (Participant 50)
○ Quote 4: “Movement stilted which made identifying slightly lame sheep virtually impossible.” (Participant 51)
○ Quote 5: “Very basic. In this game the sheep were fairly stationary which made that hard” (Participant 70)
· Sub-theme 3: Unable to mark non-lame sheep
○ Quote 1: “It was a bit frustrating not to be able to mark non-lame sheep when surveying, but that is more realistic and requires strategy.” (Participant 57)
· Sub-theme 4: Usability and Animation/simulation issues (e.g., transitions, controls, graphics) (UA)
○ Quote 1: “Game animations were not smooth, making the distinction between a normal walking gait and a limp less easy to discern.” (Participant 19)
○ Quote 2: “The graphics werent very clear - it was hard to see if they were holding a leg slightly up.” (Participant 34)
○ Quote 3: “would have enjoyed this game better if the this game if the controls worked better” (Participant 35)
○ Quote 4: “… On my PC there was a foot slide effect. I didn’t look for standing signs as I thought they were more graphics errors” (Participant 57)
3. Emotional responses to the game (ER)

· Sub-theme 1: Enjoyment
○ Quote 1: “:) [Happy face]” (Participant 15)
○ Quote 2: “: it was entertaining” [Sic] (Participant 43)
○ Quote 3: “I enjoyed the game” (Participant 44)
· Sub-theme 2: Surprise/interesting
○ Quote 1: “I thought this was brilliant” (Participant 57)
○ Quote 2: “It’s interesting to be looking for sign in virtual sheep” (Participant 68)
· Sub-theme 3: Boredom
○ Quote 1: “I got bored waiting for the sheep to move unfortunately” (Participant 8)
○ Quote 2: “I got bored waiting for them to walk” (Participant 18)
· Sub-theme 4: Frustration
○ Quote 1: “i got annoyed waiting for the sheep to move” [Sic] (Participant 27)
○ Quote 2: “Found it very frustrating.” (Participant 51)
○ Quote 3: “But I got frustrated.” (Participant 68)
· Sub-theme 5: Lack of appeal
○ Quote 1: “This sort of game doesn’t appeal to me I’m afraid. I’ve always worked in the real world.” (Participant 36)
4. Reflective experiences

○ Quote 1: “Lame sheep aren’t always that easy to spot in a field” (Participant 22)
○ Quote 2: “in a flock i would walk around them and the sheep would move” (Participant 27)
○ Quote 3: “it allowed me to get a better sense of my knowledge and skills” (Participant 44) o Quote 4: “In real life I would walk around the flock and observe they way the moved” (Participant 70)
5. Participants’ suggestions for improvements

· Sub-theme 1: Making sheep move e.g., using additional mechanisms and characters
○ Quote 1: “If there was a way to make each sheep move, that would really help to keep engagement” (Participant 8)
○ Quote 2: “If you wanted to complete the game in a shorter time, you would want the sheep to move around more. Needs a dog to run round them!” (Participant 18)
○ Quote 3: “Would be good to get sheep to move, maybe by walking a person around so they walk away from you.” (Participant 24)
○ Quote 4: “would be nice to have a method of encouraging sheep to move.” (Participant 70)
· Sub-theme 2: Providing additional visual/sound feedback
○ Quote 1: “i felt there could be improvements made as you chose the right animals maybe a sound so you know your going the right way or a counter in the corner.” (Participant 43)
○ Quote 2: “Could be enhanced by slightly more realistic depiction of sheep movement for non-lame sheep” (Participant 67[JM1])

### Supplementary Material 4: Project Budget Breakdown

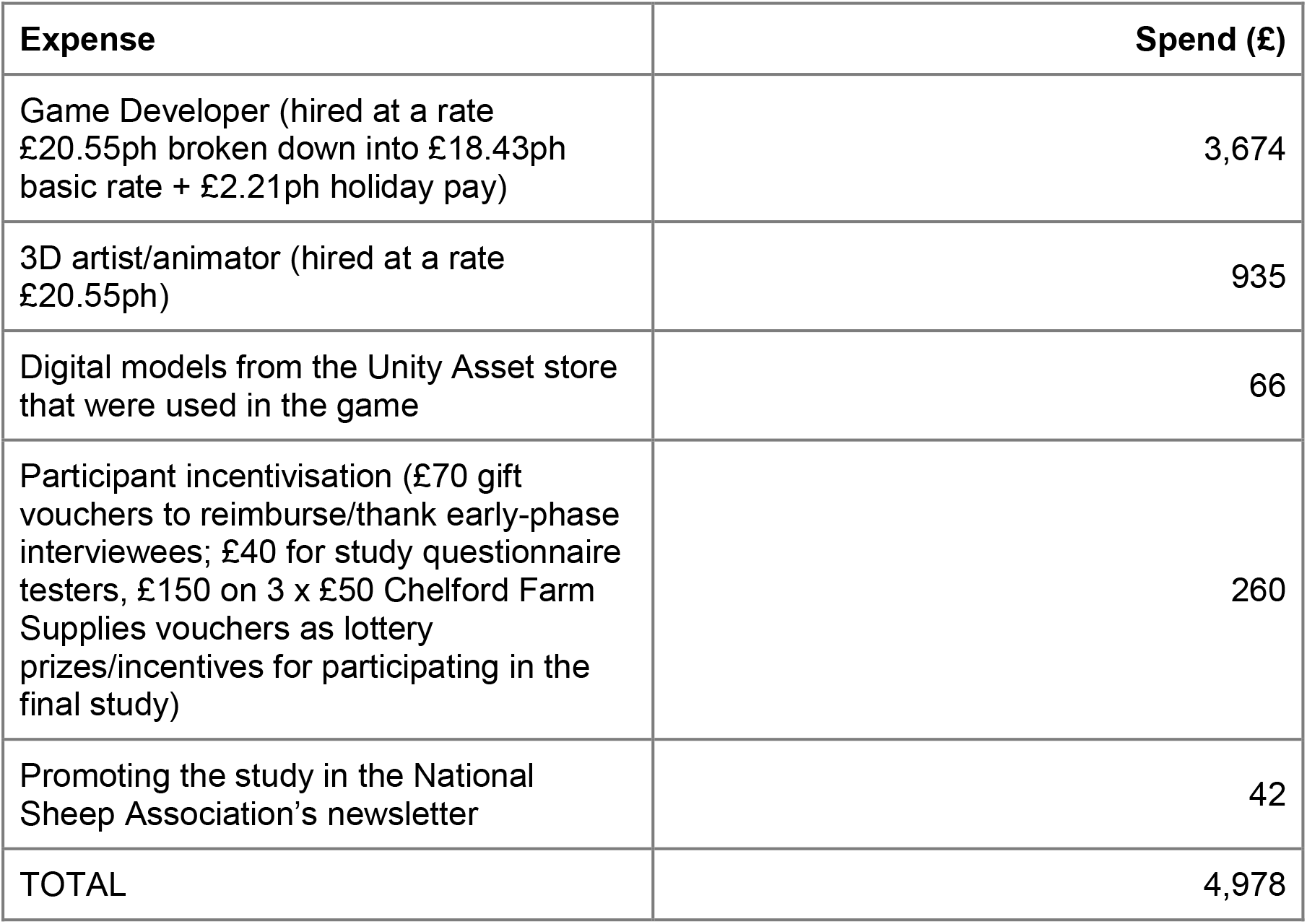

**Supplementary Figure 1:**
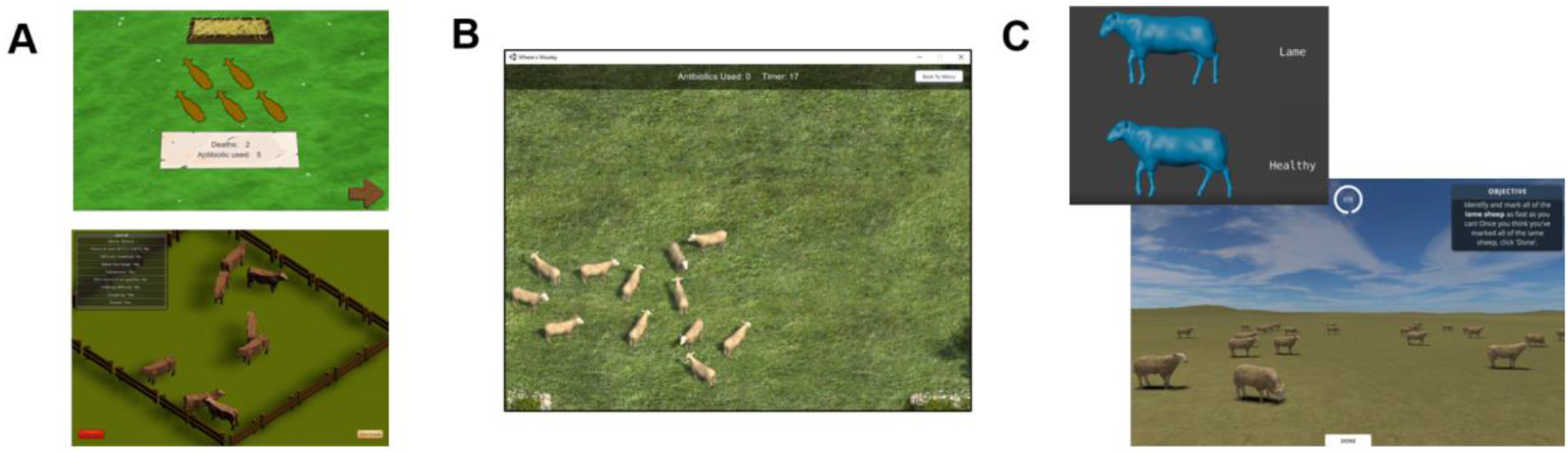
Description of the human-centered design process. A) NScreenshots of the initial wireframes/prototypes 1a and 1b developed around the concept of antibiotic use in livestock farming; B) Screenshot of prototype 2, ‘Where’s Woolly’ in which players are challenged to identify simple signs of illness in sheep; C) Screenshot of the animations and final game in which they were used, which is the game evaluated in this study.

**Supplementary Figure 2:**
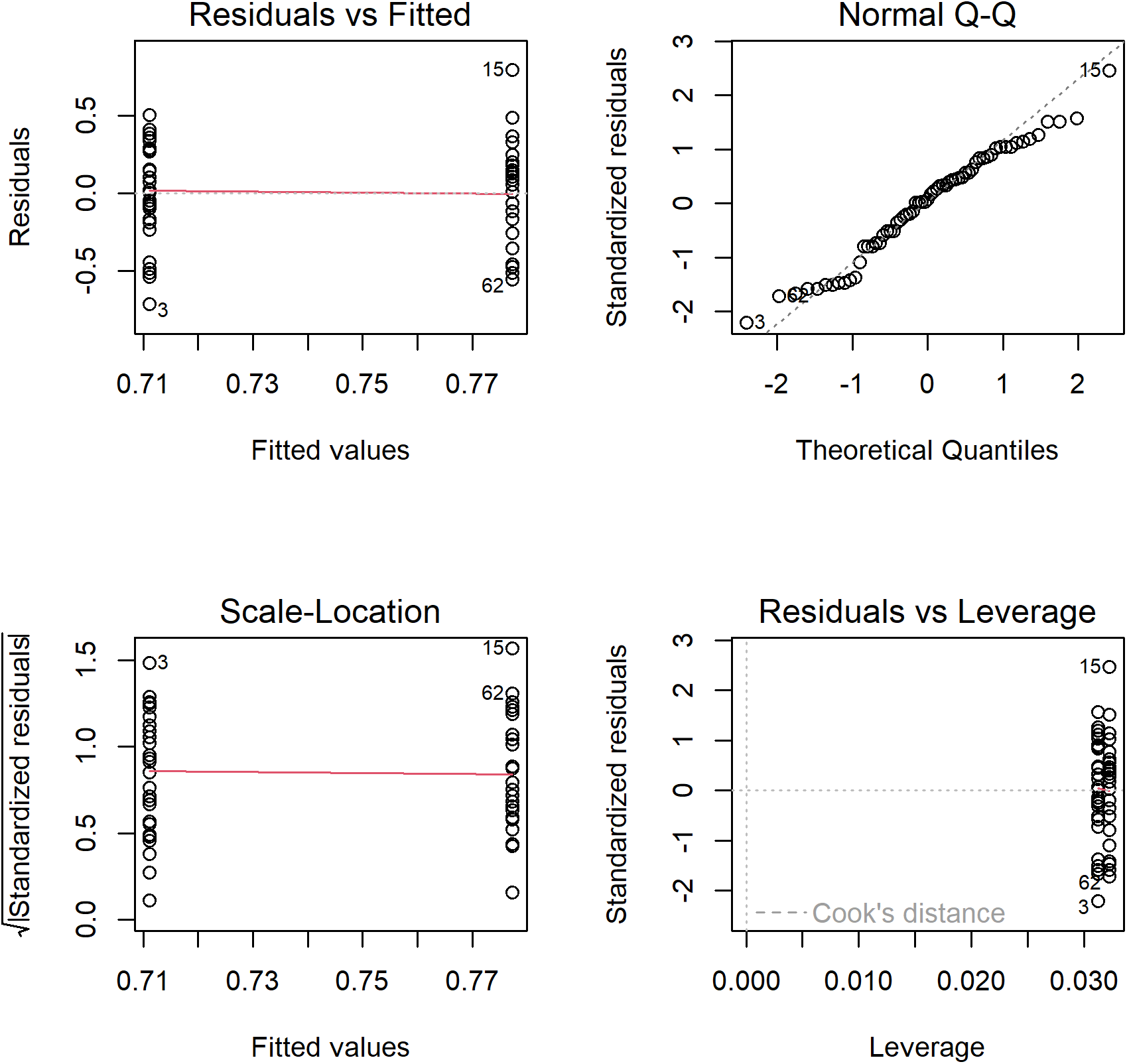
Diagnostic plots for the ‘Farming Experience’ model. From top left to bottom right: Residuals vs Fitted plot showing if residuals have linear patterns (residuals should be approximately equally spread around the horizontal red line if so); Normal Q-Q plot showing if residuals are normally distributed (residuals should approximately follow the dashed line if so); Scale-Location plot showing if residuals are spread equally along the ranges of the predictors (residuals should be approximately equally spread around the horizontal red line if so); Residuals vs Leverage plot to identify any outliers that are influential in the linear regression (Cook’s distance lines should not be visible and/or points should all be within Cook’s distance lines)

**Supplementary Figure 3:**
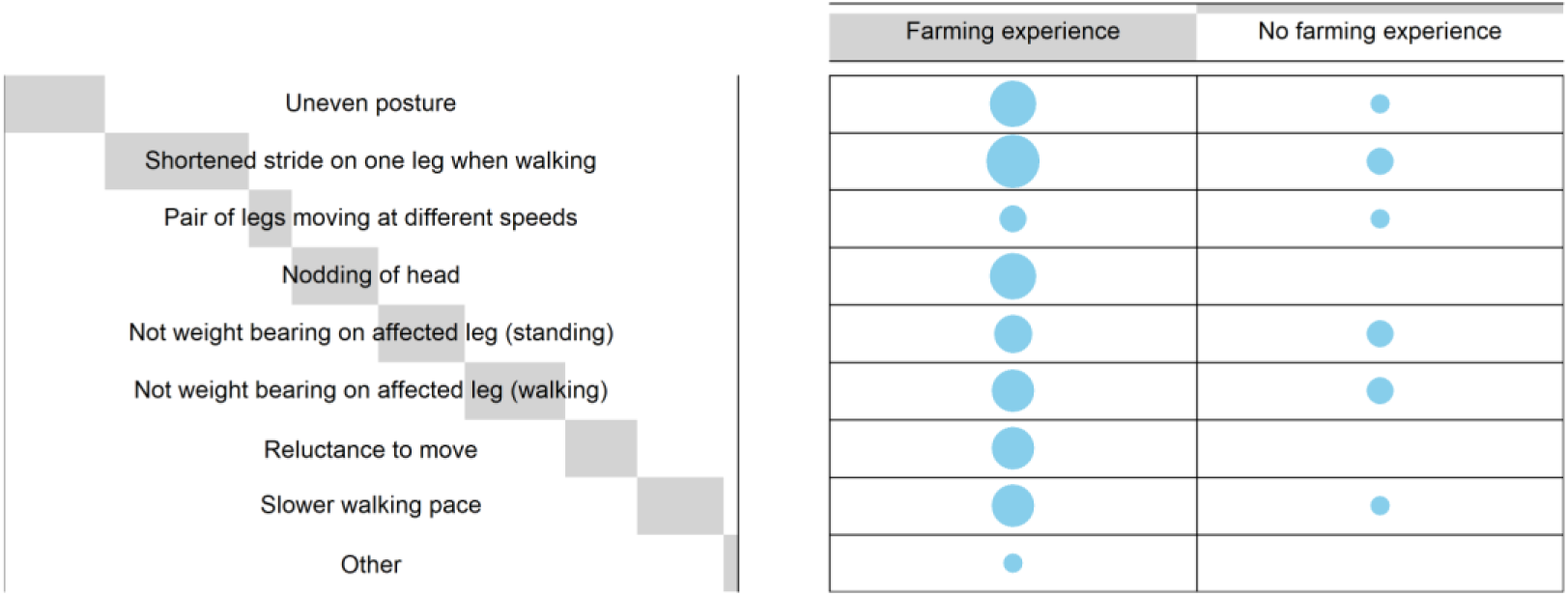
Balloon plot of contingency table used to conduct chi-squared test for a difference in the lameness signs looked for according to real-life farming experience. Size of the circles/balloons reflects the frequency of participants that looked for that lameness sign (relative to the total number of signs looked for by both groups)

**Supplementary Figure 4:**
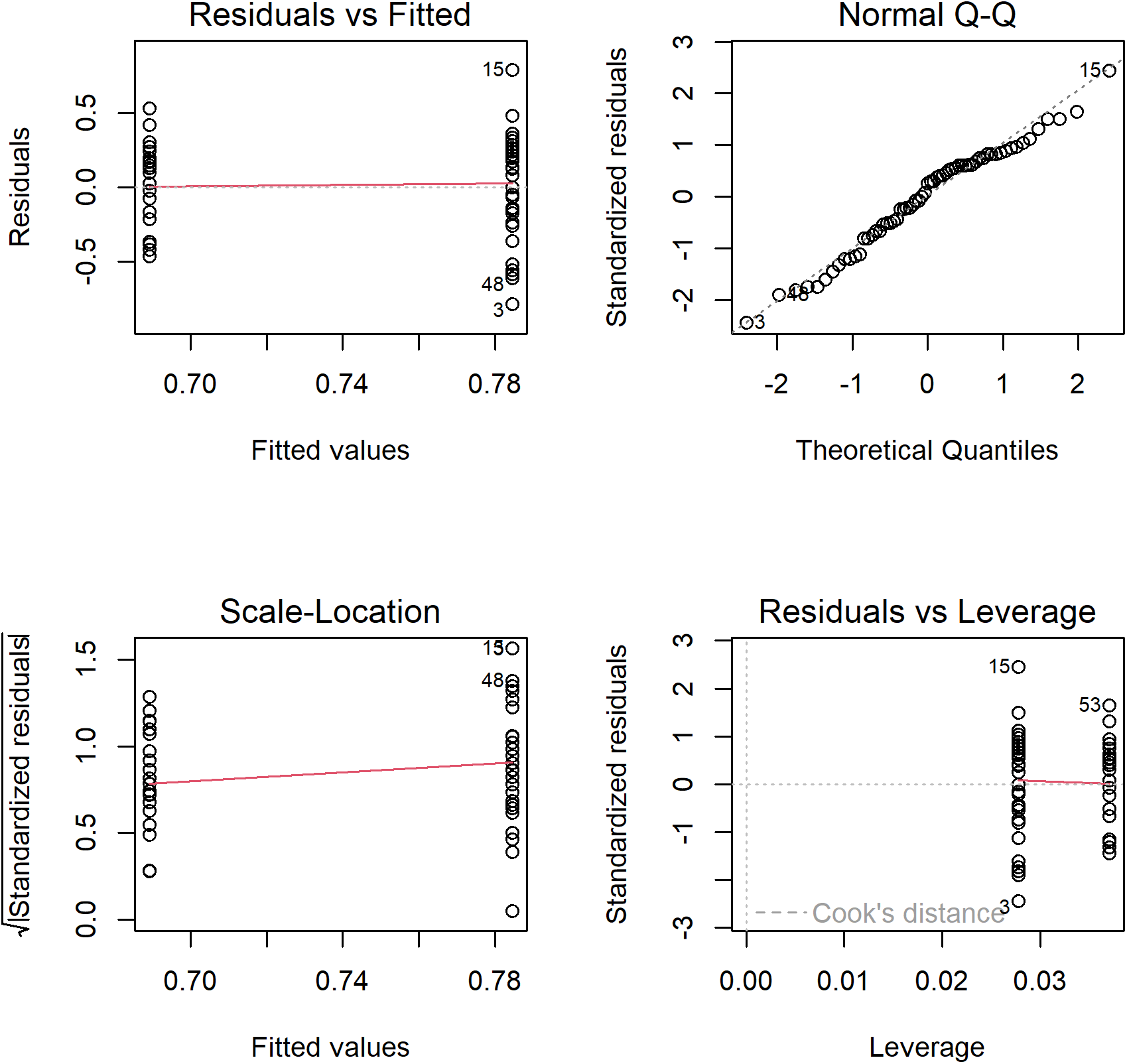
Diagnostic plots for the ‘Lameness signs looked for’ model A (uneven posture) model. From top left to bottom right: Residuals vs Fitted plot showing if residuals have linear patterns (residuals should be approximately equally spread around the horizontal red line if so); Normal Q-Q plot showing if residuals are normally distributed (residuals should approximately follow the dashed line if so); Scale-Location plot showing if residuals are spread equally along the ranges of the predictors (residuals should be approximately equally spread around the horizontal red line if so); Residuals vs Leverage plot to identify any outliers that are influential in the linear regression (Cook’s distance lines should not be visible and/or points should all be within Cook’s distance lines)

**Supplementary Figure 5:**
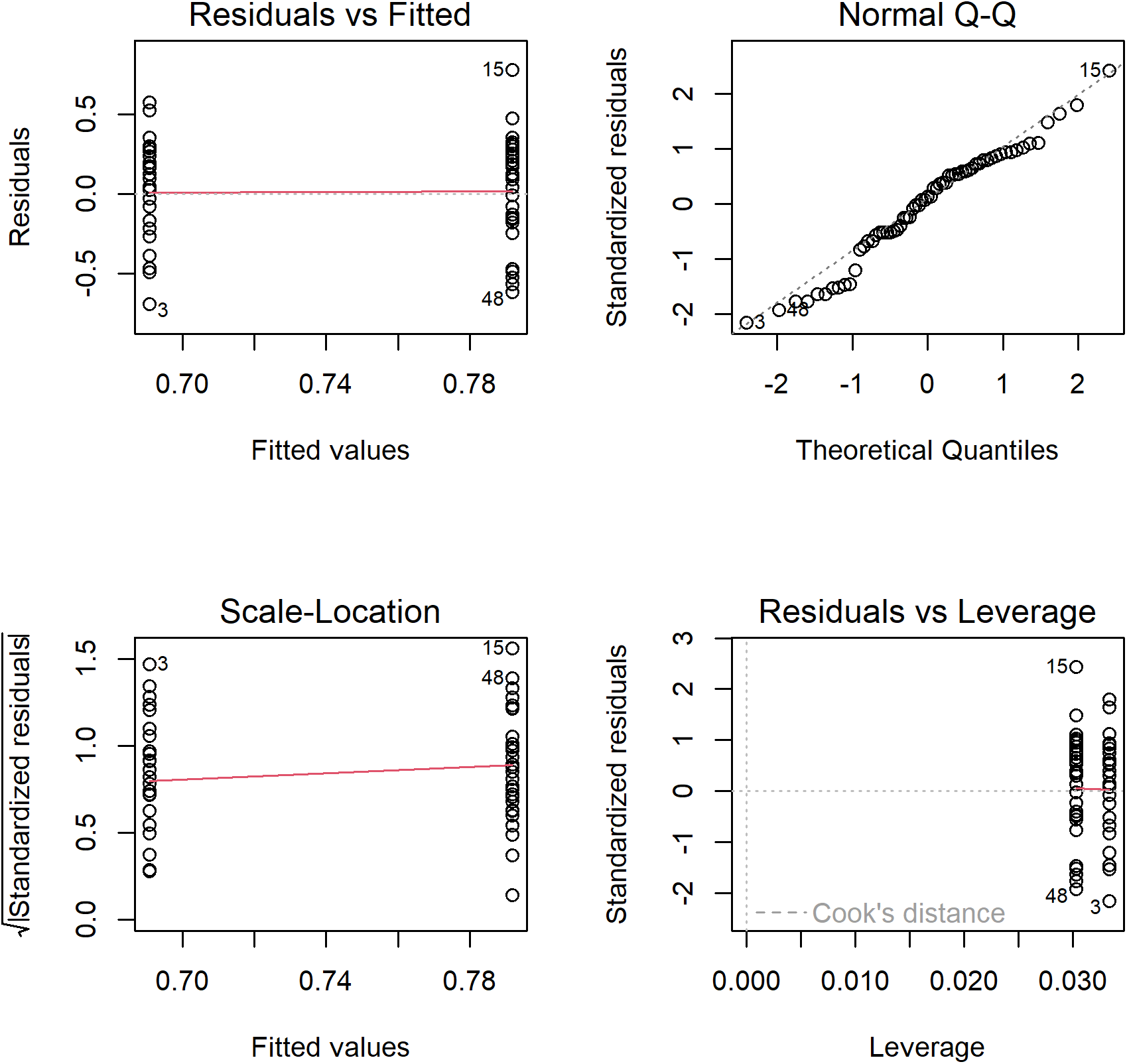
Diagnostic plots for the ‘Lameness signs looked for’ model B (limp) model. From top left to bottom right: Residuals vs Fitted plot showing if residuals have linear patterns (residuals should be approximately equally spread around the horizontal red line if so); Normal Q-Q plot showing if residuals are normally distributed (residuals should approximately follow the dashed line if so); Scale-Location plot showing if residuals are spread equally along the ranges of the predictors (residuals should be approximately equally spread around the horizontal red line if so); Residuals vs Leverage plot to identify any outliers that are influential in the linear regression (Cook’s distance lines should not be visible and/or points should all be within Cook’s distance lines)

**Supplementary Figure 6:**
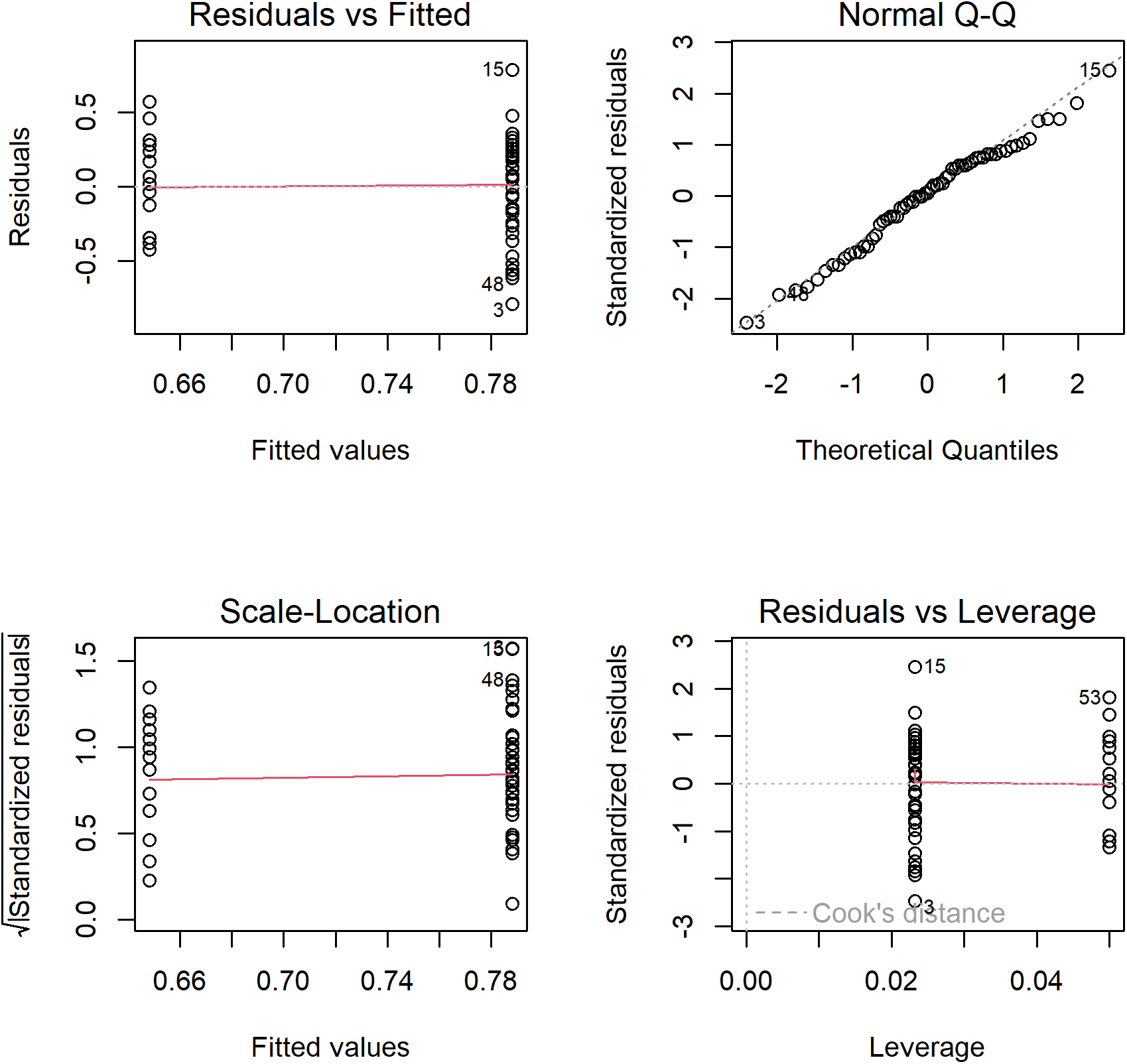
Diagnostic plots for the ‘Lameness signs looked for’ model C (raised leg) model. From top left to bottom right: Residuals vs Fitted plot showing if residuals have linear patterns (residuals should be approximately equally spread around the horizontal red line if so); Normal Q-Q plot showing if residuals are normally distributed (residuals should approximately follow the dashed line if so); Scale-Location plot showing if residuals are spread equally along the ranges of the predictors (residuals should be approximately equally spread around the horizontal red line if so); Residuals vs Leverage plot to identify any outliers that are influential in the linear regression (Cook’s distance lines should not be visible and/or points should all be within Cook’s distance lines)

**Supplementary Figure 7:**
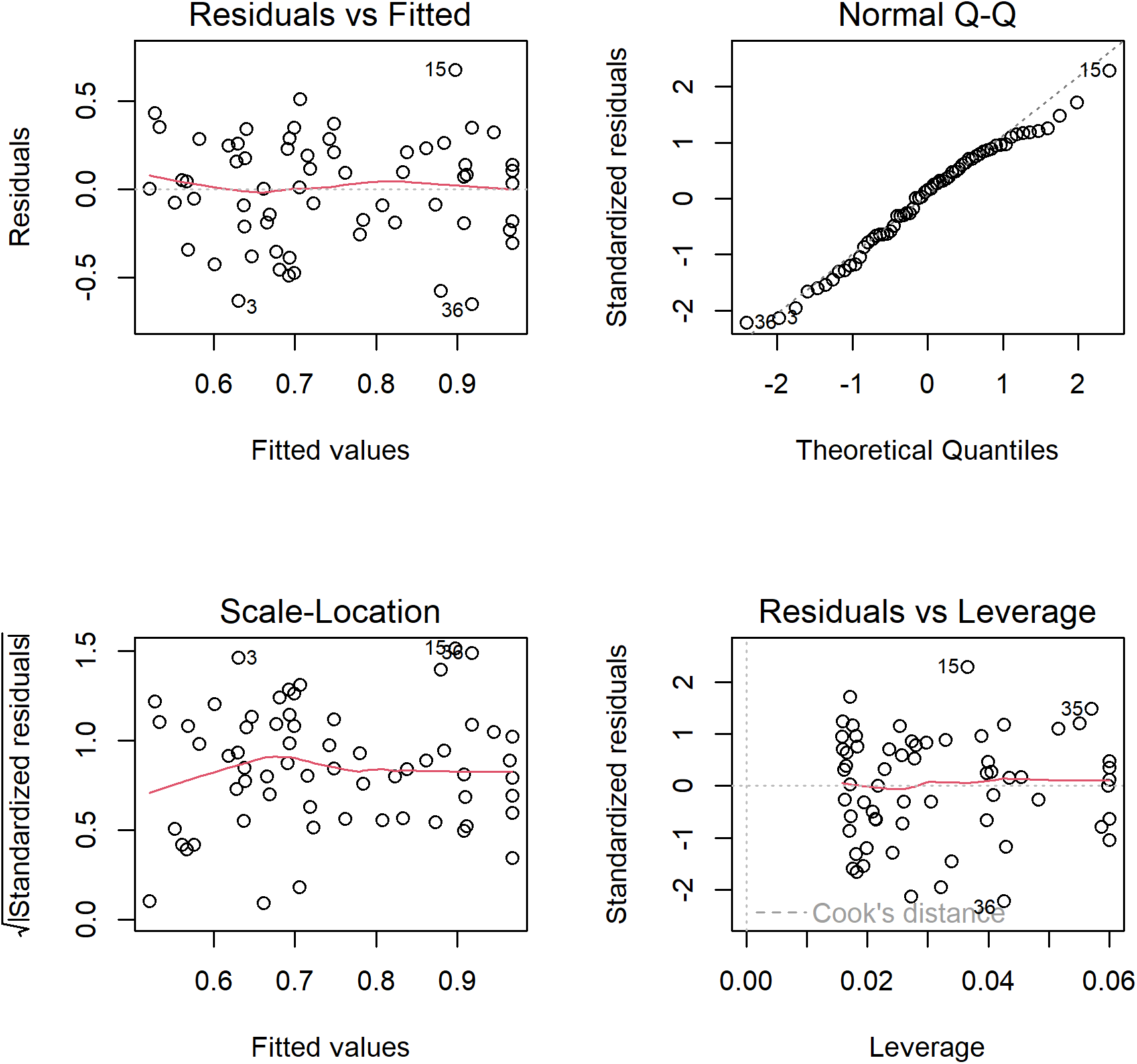
Diagnositc plots for the ‘User Engagement’ model. From top left to bottom right: Residuals vs Fitted plot showing if residuals have linear patterns (residuals should be approximately equally spread around the horizontal red line if so); Normal Q-Q plot showing if residuals are normally distributed (residuals should approximately follow the dashed line if so); Scale-Location plot showing if residuals are spread equally along the ranges of the predictors (residuals should be approximately equally spread around the horizontal red line if so); Residuals vs Leverage plot to identify any outliers that are influential in the linear regression (Cook’s distance lines should not be visible and/or points should all be within Cook’s distance lines)

## References

Barber, Stuart. 2016. Development of 4D Farms to Improve Student Learning and Safety: Final Report 2016.

Berthet, Elsa T. A., Cécile Barnaud, Nathalie Girard, Julie Labatut, and Guillaume Martin. 2016. “How to Foster Agroecological Innovations? A Comparison of Participatory Design Methods.” Journal of Environmental Planning and Management 59 (2): 280–301. https://doi.org/10.1080/09640568.2015.1009627.

Best, Caroline M., Alison Z. Pyatt, Janet Roden, Malgorzata Behnke, and Kate Phillips. 2021. “Sheep Farmers’ Attitudes Towards Lameness Control: Qualitative Exploration of Factors Affecting Adoption of the Lameness Five-Point Plan.” PLOS ONE 16 (2): e0246798. https://doi.org/10.1371/journal.pone.0246798.

Best, Caroline M., Janet Roden, Alison Z. Pyatt, Malgorzata Behnke, and Kate Phillips. 2020. “Uptake of the Lameness Five-Point Plan and Its Association with Farmer-Reported Lameness Prevalence: A Cross-Sectional Study of 532 UK Sheep Farmers.” Preventive Veterinary Medicine 181 (August): 105064. https://doi.org/10.1016/j.prevetmed.2020.105064.

Bicameral Studios. 2018. “Free Island Collection 3D Landscapes Unity Asset Store.” https://assetstore.unity.com/packages/3d/environments/landscapes/free-island-collection-104753.

Blender Foundation. 2021. “Blender.” https://www.blender.org/.

Bødker, S. 2000. “Scenarios in User-Centred Design—Setting the Stage for Reflection and Action.” Interacting with Computers 13 (1): 61–75. https://doi.org/10.1016/S0953-5438(00)00024-2.

Braun, Virginia, and Victoria Clarke. 2006. “Using Thematic Analysis in Psychology.” Qualitative Research in Psychology 3 (2): 77–101. https://doi.org/10.1191/1478088706qp063oa.

Braun, Virginia, and Victoria Clarke. 2006. 2021a. “Can I Use TA? Should I Use TA? Should I Not Use TA? Comparing Reflexive Thematic Analysis and Other Pattern-Based Qualitative Analytic Approaches.” Counselling and Psychotherapy Research 21 (1): 37–47. https://doi.org/10.1002/capr.12360.

Braun, Virginia, and Victoria Clarke. 2006. 2021b. Thematic Analysis: A Practical Guide. SAGE Publications Ltd. https://uk.sagepub.com/en-gb/eur/thematic-analysis/book248481.

Bueno, L., and Y. Ruckebusch. 1979. “Ingestive Behaviour in Sheep Under Field Conditions.” Applied Animal Ethology 5 (2): 179–87. https://doi.org/10.1016/0304-3762(79)90089-0.

Champely, Stephane, Claus Ekstrom, Peter Dalgaard, Jeffrey Gill, Stephan Weibelzahl, Aditya Anandkumar, Clay Ford, Robert Volcic, and Helios De Rosario. 2020. Pwr: Basic Functions for Power Analysis. https://CRAN.R-project.org/package=pwr.

Cohen, Jacob. 1977. Statistical Power Analysis for the Behavioral Sciences. Academic Press.

Crowley, Edward J., Matthew J. Silk, and Sarah L. Crowley. 2021. “The Educational Value of Virtual Ecologies in Red Dead Redemption 2.” People and Nature n/a (n/a). https://doi.org/10.1002/pan3.10242.

Davies, Peers, John G Remnant, Martin J Green, Emily Gascoigne, Nick Gibbon, Robert Hyde, Jack R Porteous, Kiera Schubert, Fiona Lovatt, and Alexander Corbishley. 2017. “Quantitative Analysis of Antibiotic Usage in British Sheep Flocks.” Veterinary Record 181 (19): 511–11. https://doi.org/10.1136/vr.104501.

Eklund, Aron, and James Trimble. 2021. Beeswarm: The Bee Swarm Plot, an Alternative to Stripchart. https://CRAN.R-project.org/package=beeswarm.

Fallman, Daniel. 2008. “The Interaction Design Research Triangle of Design Practice, Design Studies, and Design Exploration.” Design Issues 24 (3): 4–18. https://www.jstor.org/stable/25224179.

FAWC. 2011. “FAWC Opinion on Sheep Lameness.” https://www.gov.uk/government/publications/fawc-opinion-on-sheep-lameness.

Fountas, Spyros, Tsiropoulos, Zisis, Stamatelopoulos, Panagiotis, Anastasiou, Evangelos, Hutzenlaub, Tim, Radišić, Mladen, Minic, Vladan, and Rau Patrick. 2019. “A Serious Video Game for Smart Farming Technologies.” In Digitizing Agriculture Conference Proceedings. S. l.: s. n. https://efita-org.eu/wp-content/uploads/2020/03/EFITA_Proceedings_e-book.pdf.

GATES. 2019. “Gates Smart Farming.” Text. Gates Smart Farming. https://www.gates-game.eu/en.

Green, Laura, and Rachel Clifton. 2018. “Diagnosing and Managing Footrot in Sheep: An Update.” In Practice 40 (1): 17–26. https://doi.org/10.1136/inp.j4575.

Hanington, Bruce. 2017. “Empathy, Values, and Situated Action: Sustaining People and Planet Through Human Centered Design.” In Routledge Handbook of Sustainable Design, 1st ed. Routledge. https://www.taylorfrancis.com/chapters/edit/10.4324/9781315625508-19/empathy-values-situated-action-bruce-hanington.

Hernandez-Aguilera, J. Nicolas, Max Mauerman, Alexandra Herrera, Kathryn Vasilaky, Walter Baethgen, Ana Maria Loboguerrero, Rahel Diro, Yohana Tesfamariam Tekeste, and Daniel Osgood. 2020. “Games and Fieldwork in Agriculture: A Systematic Review of the 21st Century in Economics and Social Science.” Games 11 (4): 47. https://doi.org/10.3390/g11040047.

Jones, Anna. 2022. “Just Farmers.” https://www.justfarmers.org/.

Jones, Matt, Hughes, Robert, Murray, Aimee, Verdezoto, Nervo and Barnish, Maxwell. 2020. “Exploring Antibiotic Use Practices in Livestock Production Through a Novel, Game-Based Approach.” https://gw4.ac.uk/exploring-antibiotic-use-practices-in-livestock-production-through-a-novel-game-based-approach/.

Kaler, Jasmeet, and T. R. N. George. 2011. “Why Are Sheep Lame? Temporal Associations Between Severity of Foot Lesions and Severity of Lameness in 60 Sheep.” Animal Welfare 20: 1.

Kaler, Jasmeet, and Laura Green. 2008. “Recognition of Lameness and Decisions to Catch for Inspection Among Sheep Farmers and Specialists in GB.” BMC Veterinary Research 4 (1): 41. https://doi.org/10.1186/1746-6148-4-41.

Kaler, Jasmeet, Jurgen Mitsch, Jorge A. Vázquez-Diosdado, Nicola Bollard, Tania Dottorini, and Keith A. Ellis. 2019. “Automated Detection of Lameness in Sheep Using Machine Learning Approaches: Novel Insights into Behavioural Differences Among Lame and Non-Lame Sheep.” Royal Society Open Science 7 (1): 190824. https://doi.org/10.1098/rsos.190824.

Komsta, Lukasz, and Frederick Novomestky. 2022. Moments: Moments, Cumulants, Skewness, Kurtosis and Related Tests. https://CRAN.R-project.org/package=moments.

Lane, Rick. 2018. “Meet the Real-Life Farmers Who Play Farming Simulator | Simulation Games | the Guardian.” https://www.theguardian.com/games/2018/jul/24/meet-the-real-life-farmers-who-play-farming-simulator.

Lehtonen, Sami. 2017. “sFuture Targeting 3D Props Unity Asset Store.” https://assetstore.unity.com/packages/3d/props/sfuture-targeting-83113.

Lim, Youn-Kyung, Erik Stolterman, and Josh Tenenberg. 2008. “The Anatomy of Prototypes: Prototypes as Filters, Prototypes as Manifestations of Design Ideas.” ACM Transactions on Computer-Human Interaction 15 (2): 7:1–27. https://doi.org/10.1145/1375761.1375762.

Michsky. 2021. “Modern UI Pack GUI Tools Unity Asset Store.” https://assetstore.unity.com/packages/tools/gui/modern-ui-pack-201717.

Monk, Andrew F. 2002. “Fun, Communication and Dependability: Extending the Concept of Usability.” In People and Computers XVI - Memorable Yet Invisible, edited by Xristine Faulkner, Janet Finlay, and Françoise Détienne, 3–14. London: Springer. https://doi.org/10.1007/978-1-4471-0105-5_1.

Moojen, Fernanda Gomes, Paulo César de Faccio Carvalho, Davi Teixeira dos Santos, Armindo Barth Neto, Paulo Cardozo Vieira, and Julie Ryschawy. 2022. “A Serious Game to Design Integrated Crop-Livestock System and Facilitate Change in Mindset Toward System Thinking.” Agronomy for Sustainable Development 42 (3): 35. https://doi.org/10.1007/s13593-022-00777-5.

Nalon, Elena, and Peter Stevenson. 2019. “Addressing Lameness in Farmed Animals: An Urgent Need to Achieve Compliance with EU Animal Welfare Law.” Animals 9 (8): 576. https://doi.org/10.3390/ani9080576.

Nieuwhof, G. J., and S. C. Bishop. 2005. “Costs of the Major Endemic Diseases of Sheep in Great Britain and the Potential Benefits of Reduction in Disease Impact.” Animal Science 81 (1): 23–29. https://doi.org/10.1079/ASC41010023.

Nuritha, Ifrina, Vandha Pradwiyasma Widartha, and Saiful Bukhori. 2017. “Designing Gamification on Social Agriculture (SociAg) Application to Increase End-User Engagement.” In 2017 4th International Conference on Computer Applications and Information Processing Technology (CAIPT), 1–5. https://doi.org/10.1109/CAIPT.2017.8320713.

Pavlenko, T., D.s. Paraforos, D. Fenrich, S., A. Murdoch, R. Tranter, Y. Gadanakis, M. Arnoult, and T. Engel. 2021. “96. Increasing Adoption of Precision Agriculture via Gamification: The Farming Simulator Case.” In Precision Agriculture ?21, 803–10. Wageningen Academic Publishers. https://doi.org/10.3920/978-90-8686-916-9_96.

Petri, Giani, Christiane Gresse von Wangenheim, and Adriano Ferreti Borgatto. 2017. “MEEGA+, Systematic Model to Evaluate Educational Games.” In, edited by Newton Lee, 1–7. Cham: Springer International Publishing. https://doi.org/10.1007/978-3-319-08234-9_214-1.

Prosser, Naomi S., Kevin J. Purdy, and Laura E. Green. 2019. “Increase in the Flock Prevalence of Lameness in Ewes Is Associated with a Reduction in Farmers Using Evidence-Based Management of Prompt Treatment: A Longitudinal Observational Study of 154 English Sheep Flocks 20132015.” Preventive Veterinary Medicine 173 (December): 104801. https://doi.org/10.1016/j.prevetmed.2019.104801.

R Core Team. 2017. R: A Language and Environment for Statistical Computing. Vienna, Austria: R Foundation for Statistical Computing. https://www.R-project.org/.

Red Deer. 2020. “Sheep Realistic Characters Unity Asset Store.” https://assetstore.unity.com/packages/3d/characters/animals/mammals/sheep-realistic-176904.

RStudio Team. 2020. “RStudio: Integrated Development Environment for R.” Boston, MA. http://www.rstudio.com/.

Sutherland, Lee-Ann. 2020. “The ‘Desk-Chair Countryside’: Affect, Authenticity and the Rural Idyll in a Farming Computer Game.” Journal of Rural Studies 78 (August): 350–63. https://doi.org/10.1016/j.jrurstud.2020.05.002.

Szilágyi, Robert, Tamás Kovács, Krisztián Nagy, and László Várallyai. 2017. “Development of Farm Simulation Application, an Example for Gamification in Higher Education.” Journal of Agricultural Informatics 8 (August). https://doi.org/10.17700/jai.2017.8.2.373.

Treiblmaier, Horst, Lisa-maria Putz, and Paul Benjamin Lowry. 2018. “Setting a Definition, Context, and Theory-Based Research Agenda for the Gamification of Non-Gaming Applications.” https://papers.ssrn.com/abstract=3202034.

Tukey, John W. 1977. Exploratory Data Analysis. Vol. 2. Reading, MA.

Unity Technologies. 2021. “Unity.” https://unity.com/.

Wassink, G. J., E. M. King, R. Grogono-Thomas, J. C. Brown, L. J. Moore, and L. E. Green. 2010. “A Within Farm Clinical Trial to Compare Two Treatments (Parenteral Antibacterials and Hoof Trimming) for Sheep Lame with Footrot.” Preventive Veterinary Medicine 96 (1): 93–103. https://doi.org/10.1016/j.prevetmed.2010.05.006.

Whay, H. R., D. C. J. Main, L. E. Green, and A. J. F. Webster. 2003. “Assessment of the Welfare of Dairy Caftle Using Animal-Based Measurements: Direct Observations and Investigation of Farm Records.” Veterinary Record 153 (7): 197–202. https://doi.org/10.1136/vr.153.7.197.

Winter, Joanne R., and Laura E. Green. 2017. “Costbenefit Analysis of Management Practices for Ewes Lame with Footrot.” The Veterinary Journal 220 (February): 1–6. https://doi.org/10.1016/j.tvjl.2016.11.010.

Winter, Joanne R., Jasmeet Kaler, Eamonn Ferguson, Amy L. KilBride, and Laura E. Green. 2015. “Changes in Prevalence of, and Risk Factors for, Lameness in Random Samples of English Sheep Flocks: 20042013.” Preventive Veterinary Medicine 122 (1): 121–28. https://doi.org/10.1016/j.prevetmed.2015.09.014.

Yoo, Hwan-Soo, and Seong-Whan Kim. 2014. “Virtual Farmers Training: Realistic Simulation with Amusements Using Historic Simulation and Game Storyline.” International Journal of Multimedia and Ubiquitous Engineering 9 (5): 121–30. https://doi.org/10.14257/ijmue.2014.9.5.11.

